# Arc/Arg3.1 binds the nuclear polyadenylate-binding protein RRM and regulates neuronal activity-dependent formation of nuclear speckles

**DOI:** 10.1101/2024.09.04.610023

**Authors:** Tambudzai Kanhema, Kamil Parobczak, Sudarshan Patil, Dagmara Holm-Kaczmarek, Erik Hallin, Jan Ludwiczak, Andrzej A. Szczepankiewicz, Francois P. Pauzin, Adrian Szum, Yuta Ishizuka, Stanisław Dunin-Horkawicz, Petri Kursula, Grzegorz Wilczynski, Adriana Magalska, Clive R. Bramham

## Abstract

Arc is a neuronal activity-induced protein interaction hub with critical roles in synaptic plasticity and memory. Arc localizes to synapses and the nucleus, but its nuclear functions are little known. We show that Arc accumulates in the interchromatin space of dentate granule cell nuclei and the nucleosol subcellular fraction following seizure activity and in vivo dentate gyrus LTP. Proteomic analysis of affinity-purified Arc complexes identified proteins with functions in post-transcriptional mRNA processing. During LTP, Arc undergoes enhanced complex formation with polyadenylate binding protein nuclear 1 (PABPN1) and paraspeckle splicing factor (PSF) in the nucleosol. In vitro peptide binding arrays show selective binding of Arc to the PABPN1 polyA RNA recognition motif. In hippocampal neuronal cultures, Arc knockdown increases formation of PABPN1 nuclear speckles and blocks chemical-LTP associated increases in small PABPN1 foci. These results implicate Arc in basal and neuronal activity-dependent regulation of PABPN1 speckles involved in mRNA processing and polyadenylation.

## Introduction

The activity-regulated cytoskeleton-associated protein (Arc), also known as Arg3.1, is an immediate early gene product with critical roles in long-term synaptic plasticity (Bramham et al., 2010; Korb and Finkbeiner, 2011; Shepherd and Bear, 2011; Zhang and Bramham, 2021), and mechanisms of postnatal cortical development and memory consolidation (Guzowski et al., 2000; Plath et al., 2006; McCurry et al., 2010; Trent et al., 2015; Yang et al., 2023). Arc is also implicated in maladaptive mechanisms in substance abuse disorders (Salery et al., 2016; Pagano et al., 2023), as well as recovery from ischemic brain injury (Kraft et al., 2018; Chen et al., 2020, 2023).

Arc is transcribed as an immediate early gene in activated postsynaptic excitatory neurons, followed by transport of mRNA along dendrites to synaptic sites for local Arc synthesis. Evidence suggests that Arc functions as a protein interaction hub. In long-term depression (LTD) and synaptic scaling, Arc recruits machinery for clathrin-mediated endocytosis of AMPA-type glutamate receptors (Chowdhury et al., 2006; Shepherd et al., 2006; Okuno et al., 2012; DaSilva et al., 2016). In long-term potentiation (LTP), Arc interacts with drebrin A and promotes stabilization of actin filaments (Messaoudi et al., 2007; Nair et al., 2017), important for structural enlargement of dendritic spines (Fukazawa et al., 2003; Peebles et al., 2010; Bosch et al., 2014).

While synaptic mechanisms have received the majority of attention, Arc also accumulates in the nucleus (Bloomer et al., 2007; Korb et al., 2013; Steward et al., 2015), where it interacts with a nuclear spectrin isoform (SpIV5), and associates with promyelocytic leukemia (PML) bodies (Bloomer et al., 2007; Korb et al., 2013). Arc association with PML bodies is coupled to downregulation of AMPA receptor GluA1 transcription and homeostatic downscaling of excitatory transmission (Korb et al., 2013). Evidence for ectopic expression of Arc also shows complex formation with the histone deacetylase TIP60 involved in regulation of chromatin structure (Bloomer et al., 2007; Korb et al., 2013; Wee et al., 2014; Oey et al., 2015) However, binary protein-protein interactions and causal roles of Arc in the nucleus remained to be defined. Here, we sought to identify nuclear Arc binding partners in the adult rat brain in vivo. Using hippocampal seizure models and LTP induction in the perforant path input to the dentate gyrus (DG), we show that Arc accumulates in the interchromatin space of dentate granule cells and the nucleosol subcellular fraction. Proteomic mass spectrometric analysis of immunoprecipitated Arc complexes from the DG nucleosol identified proteins involved in post-transcriptional RNA processing and splicing. Co-immunoprecipitation and GST-Arc pulldown analysis show Arc complex formation with nuclear poly-A binding protein (PABPN1) and the paraspeckles protein, PSF (polypyrimidine tract-binding protein-associated splicing factor). This complex is detected under basal conditions and massively enhanced following LTP induction. In vitro peptide binding arrays demonstrate binary interactions, with direct binding of Arc to the polyA RNA recognition motif of PABPN1. Functionally, Arc regulates the maintenance and activity-dependent formation of small PABPN1 nuclear speckles in hippocampal neurons. The work suggests a specialized role for Arc in regulating neuronal PABPN1 dynamics and function.

## Material and Methods

### Animals

Adult male Wistar rats (3 months old, weight 170-250 g) were obtained from Mossakowski Medical Research Center, Polish Academy of Sciences and maintained at Nencki Institute Animal House. Wistar P0 pups were acquired from the same source and used for preparation of primary neuronal cultures.

### Drug-induced seizures

Seizures were evoked by intraperitoneal injection of the proconvulsant pentylenetetrazol (PTZ) Rats received a single injection of PTZ (50 mg/kg i.p.) and were killed at 30 min, 60 min, and 120 min after injection. All animals in the experiment had general seizures between grade 3 and 5 on the modified Racine scale for behavioral grading of seizure activity (0-no seizures, 1-“wet-dog” shaking and/or increased activity, 2-limb shaking/stretching and increased activity, 3-single (1-3) clonic seizures, 4-full clonic seizures, lasting for several minutes, 5-strong and long-lasting tonic seizures and long series of clonic seizures, 6-death in result of seizures). Animals which did not develop seizures within 15 min after injection of proconvulsant were excluded from further experiments. Procedures were performed with the consent of the 1st Local Ethical Committee in Warsaw (Permission numbers LKE 306/2017, LKE 774/2015 and LKE 295/217).

### In vivo synaptic electrophysiology and LTP

In vivo electrophysiology experiments were conducted on 96 adults (60–80 days old) male Sprague-Dawley rats outbred strain (Taconic Europe, Ejby, Denmark), weighing 250-400 g. Dentate gyrus tissue was obtained from anesthetized rats. Rats had free access to food and water and were on a 12-h light/dark cycle. All experimental procedures approved by Norwegian National Research Ethics Committee in compliance with EU Directive 2010/63/EU, ARRIVE guidelines. Experiments were conducted by Federation of Laboratory and Animal Science Associations (FELASA) C course-trained and certified researchers. Rats were anesthetized using urethane (1.5 g/kg, i.p.) and fixed on stereotaxic frames as described in previous works. Briefly, medial perforant path fibers were stimulated using a bipolar electrode (NE-200, 0.5 mm tip separation, Rhodes Medical Instruments, Woodland hills, CA) placed in the angular bundle (7.9 mm posterior to the bregma, 4.2 mm lateral to the midline). Evoked field potentials were recorded by an insulated tungsten (0.075 mm; A-M Systems #7960) electrode placed in the dentate hilus (3.9 mm posterior to bregma, 2.3 mm lateral to midline). Test-pulse stimulation (0.033 Hz) was given during 20 min baseline recording before HFS and during post-HFS recording. HFS was delivered in three sessions of HFS with a 10,5 min interval and each session consisted of four, 400 Hz stimulus trains (8 pulses/train) with 10 s interval between trains. After completion of recordings, rats were decapitated, and brains were taken out and dissected on ice. Ipsilateral (treated) and contralateral DG were rapidly and carefully micro-dissected on an ice-cold plate and frozen until further use. Signals from the dentate hilus were amplified, filtered (1 Hz to 10 kHz), and digitized (25 kHz). Evoked field potentials were acquired and analyzed using DataWave Technologies (Longmont, CO) WorkBench software. The maximum fEPSP slope was measured from the leading positive peak. Four consecutive responses were averaged and plotted to show the time course of percentage changes in the fEPSP slope (relative to baseline). Student’s t-test was used for statistical analysis of baseline and post-HFS recordings.

### Immunofluorescence of brain sections

Animals were injected intraperitoneally with a lethal dose of sodium pentobarbital (130 mg/kg) and perfused with 4% PFA in ice-cold PBS. Collected brains were post-fixed overnight at 4° C in 4% PFA in PBS, soaked in 30% sucrose, flash frozen, cryosectioned into 40 µm thick sections, and stored for further use in -20 °C in anti-freezing medium containing 30 % glycerol, 30 % ethylene glycol, 10 mM phosphate buffer pH 7.4. Immunofluorescence staining was performed according to routine procedures (Hall et al., 2016). Free floating brain sections were washed several times in 1xPBS, and blocked in 5 % Normal Donkey Serum (NDS, Jackson Immunoresearch) diluted in 0.1 % TritonX100 in 1xPBS (PBST). Incubation with primary antibodies (list and dilutions in Table 1) diluted in 5 % NDS in PBST was carried either for 1 h, at RT or overnight in 4 ° C in a humid chamber. Signals were detected with secondary antibodies conjugated with fluorophores (list and dilutions in Table 1) diluted in 5 % NDS in PBST. Samples were counterstained with Hoechst 33342 (Invitrogen, H3570) for 10 min RT, washed 3x for 5 min with PBST and mounted on glass slides using Vectashield (Vector Laboratories) and analyzed using a confocal microscope.

**Table 1.**
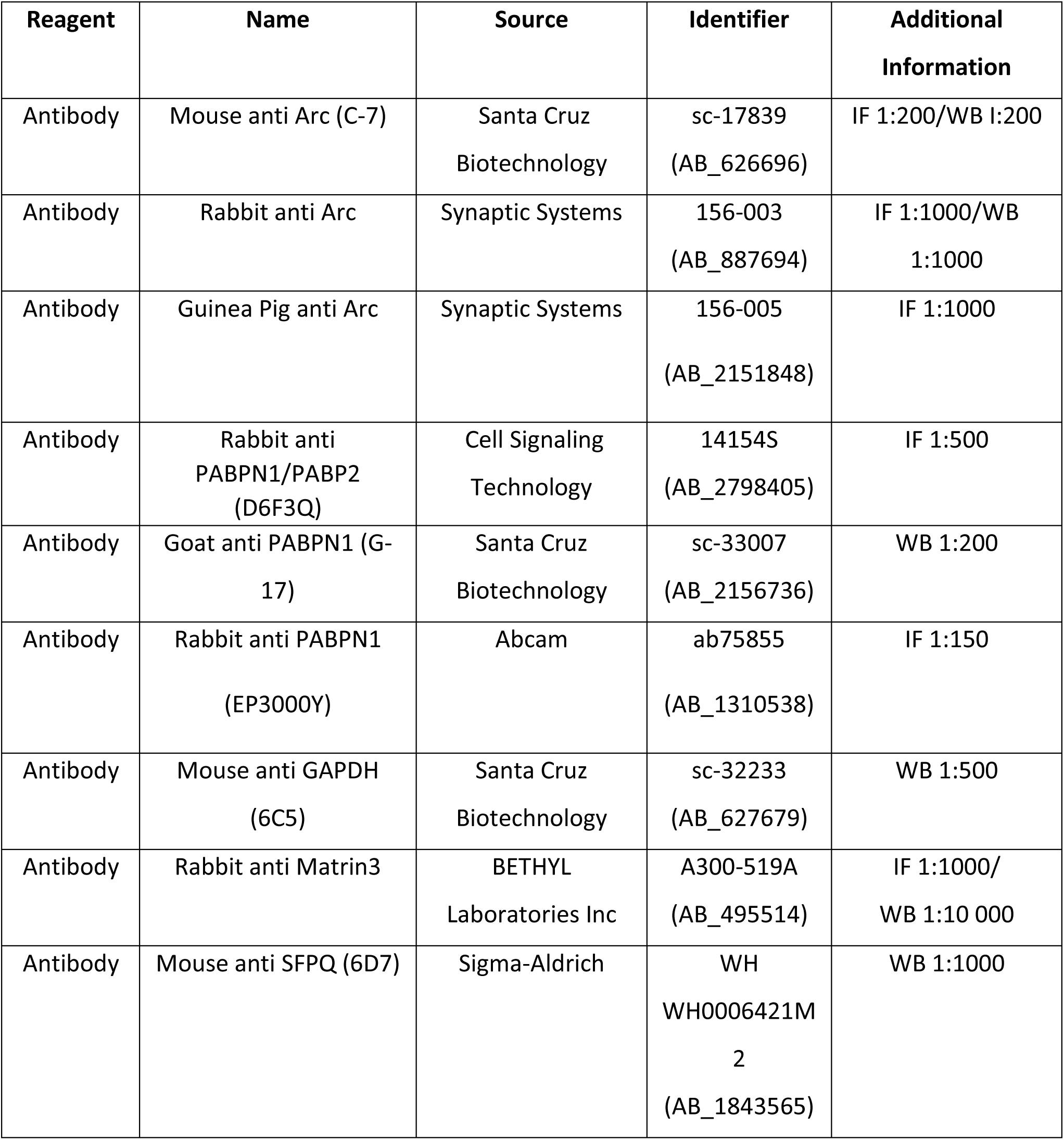
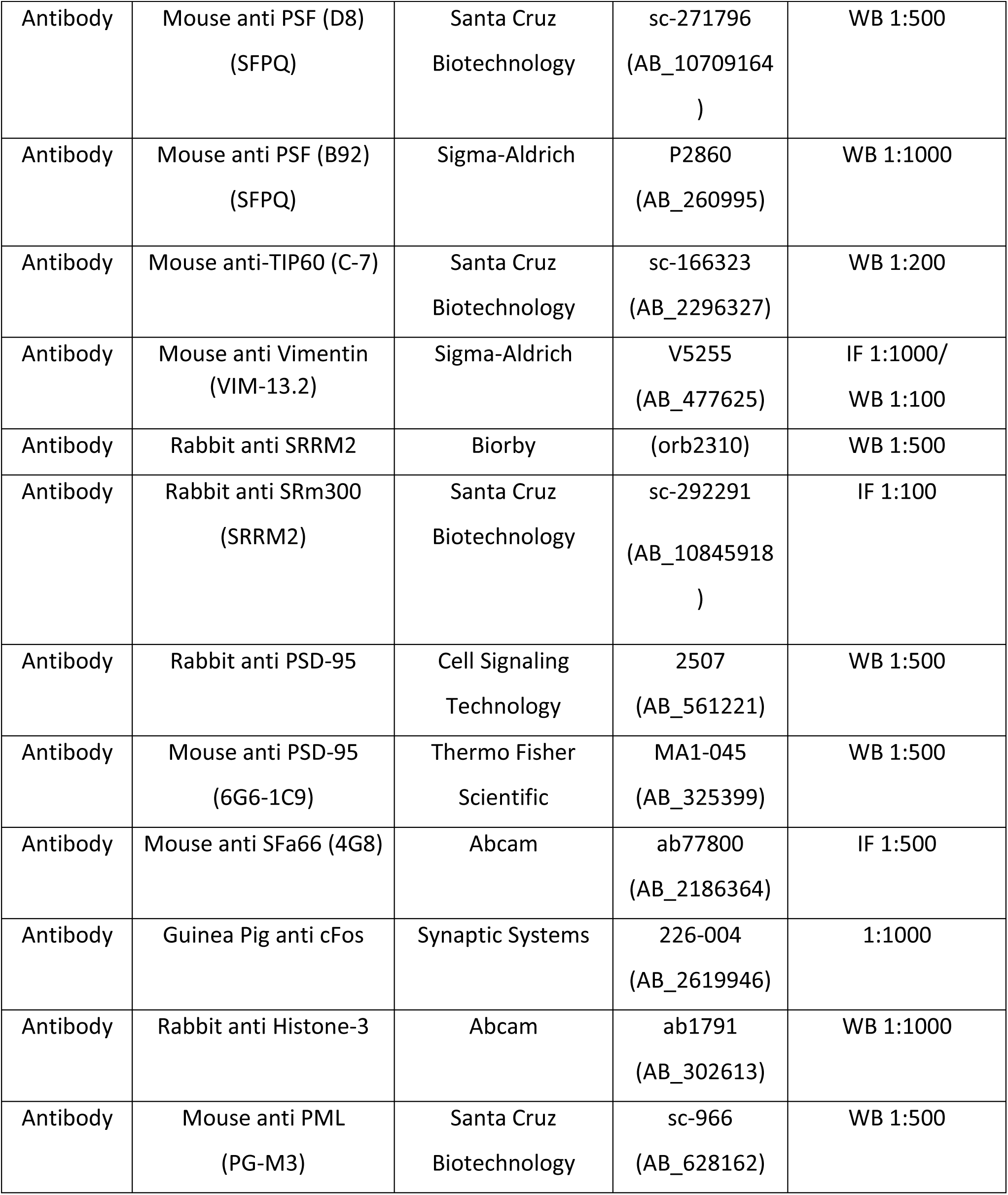
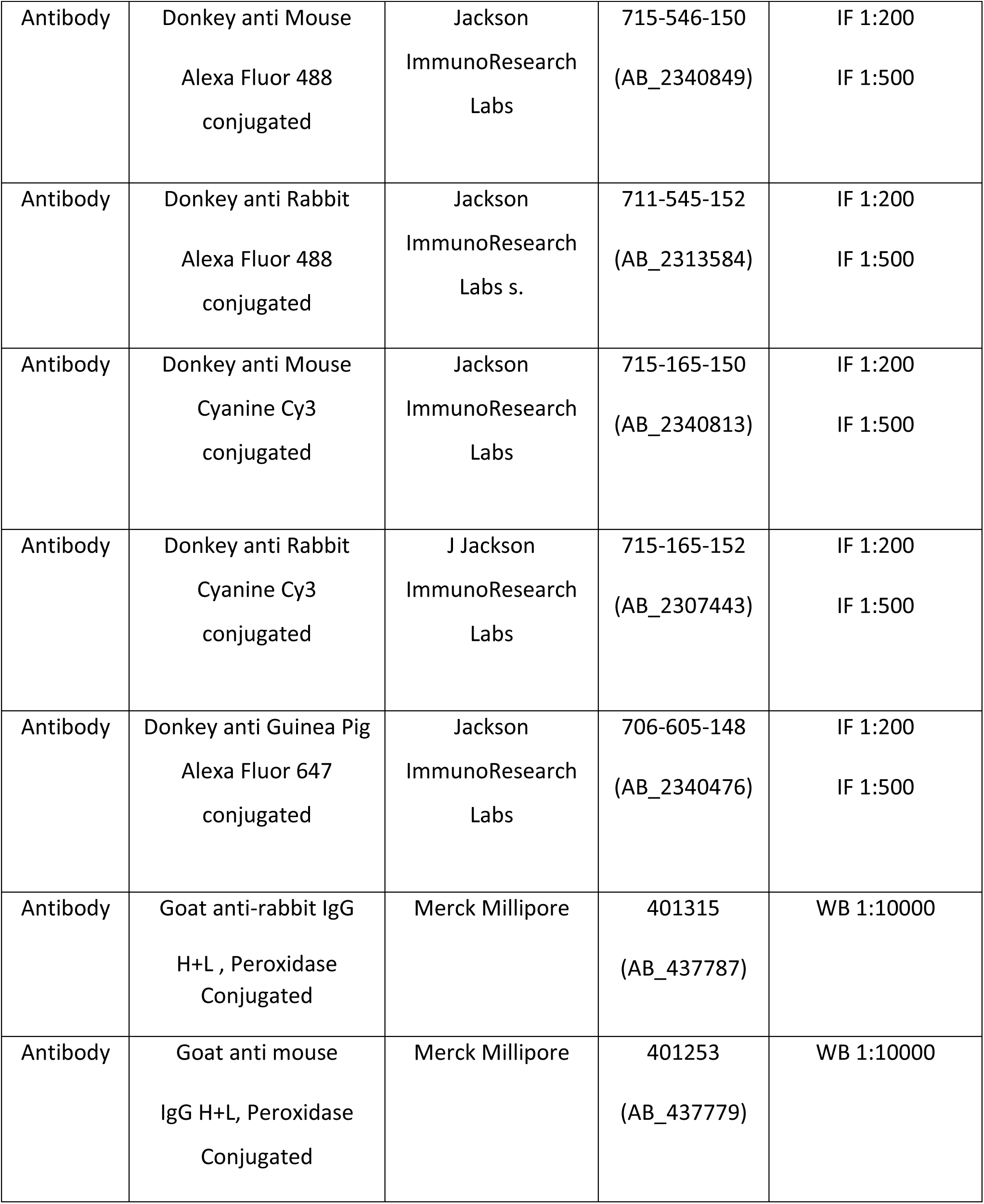
Antibodies used in this study (IF, immunofluorescence; IP, immunoprecipitation; WB, Western blot.

| Reagent | Name | Source | Identifier | Additional Information |
| --- | --- | --- | --- | --- |
| Antibody | Mouse anti Arc (C-7) | Santa Cruz<br>Biotechnology | sc-17839<br>(AB_626696) | IF 1:200/WB 1:200 |
| Antibody | Rabbit anti Arc | Synaptic Systems | 156-003<br>(AB_887694) | IF 1:1000/WB<br>1:1000 |
| Antibody | Guinea Pig anti Arc | Synaptic Systems | 156-005<br>(AB_2151848) | IF 1:1000 |
| Antibody | Rabbit anti<br>PABPN1/PABP2<br>(D6F3Q) | Cell Signaling<br>Technology | 14154S<br>(AB_2798405) | IF 1:500 |
| Antibody | Goat anti PABPN1 (G-17) | Santa Cruz<br>Biotechnology | sc-33007<br>(AB_2156736) | WB 1:200 |
| Antibody | Rabbit anti PABPN1<br>(EP3000Y) | Abcam | ab75855<br>(AB_1310538) | IF 1:150 |
| Antibody | Mouse anti GAPDH<br>(6C5) | Santa Cruz<br>Biotechnology | sc-32233<br>(AB_627679) | WB 1:500 |
| Antibody | Rabbit anti Matrin3 | BETHYL<br>Laboratories Inc | A300-519A<br>(AB_495514) | IF 1:1000/<br>WB 1:10 000 |
| Antibody | Mouse anti SFPQ (6D7) | Sigma-Aldrich | WH<br>WH0006421M<br>2<br>(AB_1843565) | WB 1:1000 |
| Antibody | Mouse anti PSF (D8)<br>(SFPQ) | Santa Cruz<br>Biotechnology | sc-271796<br>(AB_10709164<br>) | WB 1:500 |
| Antibody | Mouse anti PSF (B92)<br>(SFPQ) | Sigma-Aldrich | P2860<br>(AB_260995) | WB 1:1000 |
| Antibody | Mouse anti-TIP60 (C-7) | Santa Cruz<br>Biotechnology | sc-166323<br>(AB_2296327) | WB 1:200 |
| Antibody | Mouse anti Vimentin<br>(VIM-13.2) | Sigma-Aldrich | V5255<br>(AB_477625) | IF 1:1000/<br>WB 1:100 |
| Antibody | Rabbit anti SRRM2 | Biorby | (orb2310) | WB 1:500 |
| Antibody | Rabbit anti SRm300<br>(SRRM2) | Santa Cruz<br>Biotechnology | sc-292291<br>(AB_10845918<br>) | IF 1:100 |
| Antibody | Rabbit anti PSD-95 | Cell Signaling<br>Technology | 2507<br>(AB_561221) | WB 1:500 |
| Antibody | Mouse anti PSD-95<br>(6G6-1C9) | Thermo Fisher<br>Scientific | MA1-045<br>(AB_325399) | WB 1:500 |
| Antibody | Mouse anti SFa66 (4G8) | Abcam | ab77800<br>(AB_2186364) | IF 1:500 |
| Antibody | Guinea Pig anti cFos | Synaptic Systems | 226-004<br>(AB_2619946) | 1:1000 |
| Antibody | Rabbit anti Histone-3 | Abcam | ab1791<br>(AB_302613) | WB 1:1000 |
| Antibody | Mouse anti PML<br>(PG-M3) | Santa Cruz<br>Biotechnology | sc-966<br>(AB_628162) | WB 1:500 |
| Antibody | Donkey anti Mouse<br>Alexa Fluor 488<br>conjugated | Jackson<br>ImmunoResearch<br>Labs | 715-546-150<br>(AB_2340849) | IF 1:200<br>IF 1:500 |
| Antibody | Donkey anti Rabbit<br>Alexa Fluor 488<br>conjugated | Jackson<br>ImmunoResearch<br>Labs s. | 711-545-152<br>(AB_2313584) | IF 1:200<br>IF 1:500 |
| Antibody | Donkey anti Mouse<br>Cyanine Cy3<br>conjugated | Jackson<br>ImmunoResearch<br>Labs | 715-165-150<br>(AB_2340813) | IF 1:200<br>IF 1:500 |
| Antibody | Donkey anti Rabbit<br>Cyanine Cy3<br>conjugated | J Jackson<br>ImmunoResearch<br>Labs | 715-165-152<br>(AB_2307443) | IF 1:200<br>IF 1:500 |
| Antibody | Donkey anti Guinea Pig<br>Alexa Fluor 647<br>conjugated | Jackson<br>ImmunoResearch<br>Labs | 706-605-148<br>(AB_2340476) | IF 1:200<br>IF 1:500 |
| Antibody | Goat anti-rabbit IgG<br>H+L , Peroxidase<br>Conjugated | Merck Millipore | 401315<br>(AB_437787) | WB 1:10000 |
| Antibody | Goat anti mouse<br>IgG H+L, Peroxidase<br>Conjugated | Merck Millipore | 401253<br>(AB_437779) | WB 1:10000 |

**Table 2.**
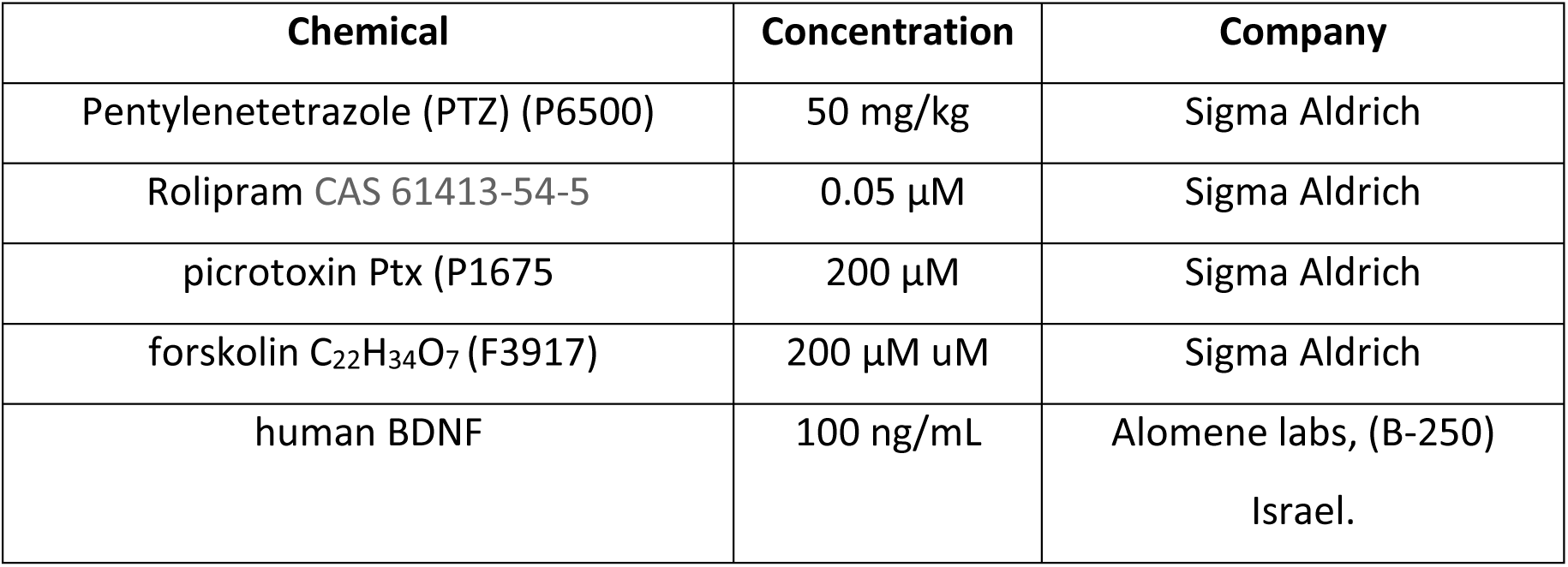
Pharmacology.

### Tissue dissection and sample preparation

At the end of electrophysiological recording, rats were decapitated, and the dentate gyri were rapidly dissected on ice. The samples were homogenized in buffer containing 50 mM Tris-HCL pH 7. 5, 150 mM KCl, 1 mM DTT, 1 mM EDTA, 1 mM PMSF, 1 % NP-40, and protease inhibitor cocktail (Roche, #11836170001). Homogenization was performed manually with 10–12 gentle strokes in a tissue grinder with a clearance of 0.1–0.15 mm (Thomas Scientific, Swedesboro, NJ). Protein concentration was measured using BCA protein assay (Thermo Fisher Scientific, # 23227). Homogenates were stored at –80 °C until use.

### Subcellular fractionation

The subcellular fractionation was performed according to (Nair et al., 2017). Rat dentate gyrus tissue was excised, weighed, and rinsed in phosphate-buffered saline (PBS) (pH 7.4). A Subcellular Protein Fractionation Kit for Tissues (Thermo Fisher Scientific # 87790) was used to separate cytoplasmic, membrane, nuclear soluble, chromatin-bound nuclear and cytoskeletal protein extracts, according to the manufacturer’s instructions. The tissue was first homogenized in the Cytoplasmic Extraction Buffer (CEB) using a tissue grinder. The homogenate was transferred to a Thermo Fisher Scientific Pierce Tissue Strainer in a 15 ml conical tube and centrifuged for 5 min at 500 x g. The strainer with debris was discarded, and the supernatant (cytoplasmic extract) was recovered. The remaining pellet was resuspended in Membrane Extraction Buffer (MEB) and incubated at 4 °C for 10 min with gently mixing. The membrane extract was recovered by centrifugation at 3000 x g for 5 min. The pellet was resuspended in Nuclear Extraction Buffer (NEB) and incubated at 4 °C for 30 min with gentle mixing. The soluble nuclear extract was separated by centrifugation at 5000 x g for 5 min. NEB containing micrococcal nuclease was added to the pellet and incubated at 37 °C for 15 min with gentle mixing. Chromatin-bound nuclear proteins were released and recovered by centrifugation of 16,000 x g for 5 min. The remaining pellet was resuspended in Pellet Extraction Buffer (PEB) and incubated at room temperature for 10 min. The cytoskeletal extract was recovered by centrifugation at 16,000 x g for 5 min.

### GST-Arc protein expression and pull-down experiments

GST-fusion proteins were expressed in BL21-Codon Plus strain of Escherichia coli cells (Agilent, Santa Clara, CA) and purified with glutathione-Sepharose beads (GE Healthcare Life Sciences, Little Chalfont, UK #1705601) according to the manufacturer’s instructions. For pull-down experiments 10 mg of unfused GST or GST-fused protein bound to glutathione-Sepharose were incubated for 1 hr at 4 °C with the nuclear extracts (1 mg of protein) in a buffer containing 50 mM Tris–HCl pH 7.4, 150 mM NaCl,1 mM EDTA, 0.5 % NP-40 supplemented with 1 mM PMSF and 1 Complete Protease Inhibitor Cocktail (Roche, Basel, Switzerland). The beads were washed three times with lysis buffer, and bound proteins were eluted by boiling in the Laemmli sample buffer and analyzed by immunoblotting.

### Co-immunoprecipitation and immunoblotting

Two different immunoprecipitation protocols were applied. The dentate gyri were either homogenized with RIPA lysis buffer (Cat R0278, Sigma) or lysis buffer containing 50mM Tris-HCl pH 7,5, 150mM KCl, 1mM DTT, 1mM EDTA, 1mM PMSF. Both buffers contained protease inhibitors (Roche, #11836170001). Tissue was homogenized using a mechanical glass-Teflon tissue homogenizer. Two µg of antibody was incubated for 30 min at RT with 20 μl of washed protein G-agarose beads or protein A/G mix magnetic beads (Cat. No. LSKMAGAG02, Millipore) or with protein G Sepharose 4 fast flow slurry (GE Healthcare #17061801) for each immunoprecipitation. 200-250 g lysate was incubated with antibody-bound beads at 4 °C for 2 hr or overnight. Immunoprecipitates were washed three times with homogenizing buffer and eluted in 2x sample buffer. Proteins were then separated by SDS-PAGE and transferred to nitrocellulose membranes and incubated with the primary antibodies.

The expression level of various proteins was assessed using western blotting. Tissue was lysed in a cold radioimmunoprecipitation assay (RIPA) buffer containing protease and phosphatase inhibitors. Tissue lysates were centrifuged at 14 000 rpm for 10 min at 4 °C. After protein isolation the protein amount measurement was done by the BSA method. Proteins in cell lysates (or immunoprecipitated samples) were resolved through SDS-PAGE. Samples were boiled in a sample buffer (Bio-Rad) and resolved via 10 % SDS-PAGE mini gels. Proteins were transferred to nitrocellulose membranes (Amersham Biosciences), blocked with 5 % nonfat dry milk, probed with horseradish peroxidase-conjugated anti-rabbit or anti-mouse secondary antibodies (1:10,000, Calbiochem) and developed using chemiluminescence reagents (ECL, Pierce). Immunoreactive bands were visualized by Gel DOC TM XRS (BIO RAD).

### Immunoprecipitation coupled mass spectrometry

The DG nucleosol fraction was used to perform Arc immunoprecipitation (IP). Each IP was done from single DG whole nucleosol minus 20 µg used as input sample. Anti-Arc antibody and IgG– Agarose (Sigma A0919) were incubated for 30 min at room temperature with 20 µl of washed protein A/G mix magnetic beads (Cat. No. LSKMAGAG02, Millipore). IgG bound beads were run as a non-immune IP control. The nuclear fraction 100-150 µg lysate was incubated with antibody-bound beads at 4 °C overnight. Immunoprecipitated samples were washed three times with PBS buffer with 0.1 % triton x 100 and proteins were eluted with SDS sample buffer and heated at 95°C for 5 min. The samples were then run on 10 % SDS-PAGE gel for 5 cm long in separation gel and stained using Bio-Safe Coomassie Stain (BioRad).

Using NanoLC-ESI-LTQ Orbitrap Elite mass spectrometry about 0.5 µg protein as tryptic peptides dissolved in 2 % acetonitrile (ACN) and 0.5 % formic acid (FA), were injected into an Ultimate 3000 RSLC system (Thermo Scientific, Sunnyvale, California, USA) connected online to a linear quadrupole ion trap-orbitrap (LTQ-Orbitrap Elite) mass spectrometer (Thermo Scientific, Bremen, Germany) equipped with a nanospray Flex ion source (Thermo Scientific). The sample was loaded and desalted on a pre-column (Acclaim PepMap 100, 2 cm x 75 µm ID nanoViper column, packed with 3µm C18 beads) at a flow rate of 6 µl/min for 5 min with 0.1 % TFA (trifluoroacetic acid, vol/vol).

Peptides were separated during a biphasic ACN gradient from two nanoflow UPLC pumps (flow rate of 200 nl /min) on a 50 cm analytical column (Acclaim PepMap 100, 25 x 75 µm ID nanoViper column, packed with 2 µm C18 beads). Solvent A and B were0.1 % FA (vol/vol) in water and 100 % ACN respectively. The gradient composition was 5 %B during trapping (5min) followed by 5-7 %B over 1 min, 7–32 %B for the next 129 min, 32-40%B over 10 min, and 40–80 %B over 5 min.

Elution of very hydrophobic peptides and conditioning of the column were performed during 20 min isocratic elution with 80%B and 20 min isocratic elution with 5 %B respectively. The eluting peptides from the LC-column were ionized in the electrospray and analyzed by the LTQ-Orbitrap Elite. The mass spectrometer was operated in the DDA-mode (data-dependent-acquisition) to automatically switch between full scan MS and MS/MS acquisition. Instrument control was through Tune 2.7.0 and Xcalibur 2.2.

Survey full scan MS spectra (from m/z 300 to 2,000) were acquired in the Orbitrap with resolution R = 240,000 at m/z 400 (after accumulation to a target value of 1e6 in the linear ion trap with maximum allowed ion accumulation time of 300ms). The 12 most intense eluting peptides above an ion threshold value of 3000 counts, and charge states 2 or higher, were sequentially isolated to a target value of 1e4 and fragmented in the high-pressure linear ion trap by low-energy CID (collision-induced-dissociation) with normalized collision energy of 35 % and wideband-activation enabled. The maximum allowed accumulation time for CID was 150 ms, with isolation width maintained at 2 Da, activation q = 0.25, and activation time of 10 ms. The resulting fragment ions were scanned out in the low-pressure ion trap at normal scan rate and recorded with the secondary electron multipliers. One MS/MS spectrum of a precursor mass was allowed before dynamic exclusion for 40 s. Lock-mass internal calibration was not enabled. The spray and ion-source parameters were as follows. Ion spray voltage = 1800 V, no sheath and auxiliary gas flow, and capillary temperature = 260 °C.

### Proteomic data analysis

Four softwares were used for this study: MaxQuant v1.5.5.1, Perseus v1.5.6.0, SPSS v25, and Excel 2019. The raw files from the LC-MS/MS were analyzed using MaxQuant version 1.5.5.1 and the integrated Andromeda. For proteins and peptides, the maximum false-discovery rate (FDR) was set to 0.01. The STRING database (STRING v.11.0; www.string-db.org) was used to identify known and predicted functional networks and to predict protein-protein interactions. Further editing of generated networks was done in Cytoscape version 3.7.2, with the use of the integrated app ClueGO.

### Densitometry and statistical analysis

Densitometric analysis was conducted using Image J Software. The blots were scanned using Gel DOC EQ (BIO RAD) and band intensities were quantified using analytical software (Quantity one 1D analysis software, BIO RAD). Densitometric values from the treated dentate gyrus (+) were expressed as fold change relative to contralateral control dentate gyrus (-). Pairwise comparisons of means were evaluated with a two-tailed Student’s t test using GraphPad prism software. The p-value for significance was 0.05.

### Peptide arrays

A custom array of partially overlapping peptides derived from the human PABPN1 sequence (NP_004634.1 and rat Arc sequence (NP_062234.1:1.0.392) was synthesized on (Celluspot Arrays, by Intavis Bioanalytical Instruments AG, Cologne, Germany). Each peptide is 14 amino acids (aa) in length with 7 aa overlapping sequences (Fig 5 F), resulting in an array covering the full-length PABPN sequence of 306 residues. The peptide array was activated in methanol, washed in Tris buffered saline Tween (TBST) (50 mM Tris-HCl (pH 7.5), 0.15 M NaCl, 0.1 % (vol/vol) (Tween 20) and blocked in TBST + 1 % BSA (wt/vol). Human recombinant GST-Arc was purified in its untagged, monomeric state as previously described (Hallin et al., 2018). The peptide array membrane was incubated in GST-Arc at a final concentration of 0.5 mg/mL overnight at 4 ⁰C. The array was washed three times for 5 min in TBST and incubated overnight at 4 ⁰C in Arc polyclonal rabbit (156-003) antibody diluted in blocking solution. The array was washed three times for 5 min with TBST and incubated for 1 hr at room temperature with secondary antibody horseradish peroxidase-conjugated anti-rabbit or anti-mouse secondary antibodies (1:10,000, Calbiochem) diluted in TBST. The array was washed three times for 5 min in TBST. The binding was detected by using Pierce ECL Western Blotting Substrate (Thermo Fisher Scientific, Waltham, MA, USA) according to the manufacturer’s instructions. Immunoreactive sequences were visualized by Gel DOC TM XRS (BIO RAD). Mapping of PABPN1 bind to Arc was done in the same way as we did when mapping Arc to PABPN1. Purified human GST-PABPN1 was obtained from GenScript.

### Structural modeling of the Arc-PABPN1 complex with experimental constraints

To gain insights into the structural basis of the ARC-PABPN1 recognition, we used the HADDOCK2 webserver (van Zundert et al., 2016) which performs protein-protein docking with additional experimental constraints. Template structures were retrieved from the PDB database – 3B4M dimer (chains A and B) for the RRM domain of human *PABPN1* (Ge et al., 2008) and 6GSE (chain A) for the capsid domain of *Rattus norvegicus Arc* (Nielsen et al., 2019). Considering that the human and rat variants of the Arc capsid domain share 97 % sequence identity, we modeled the former by substituting relevant residues using PyMOL. Since the interacting regions identified by peptide array analysis are spread across the 9-residue fragments, we performed a computational scan to indicate the residues most likely to be involved in complex formation. To this end, we performed a series of 19 docking experiments, selecting the most prominent K213 residue of one of the PABPN1 monomers and non-overlapping 3-residue fragments in region 302-358 of the Arc protein as anchor points (i.e., points expected to be in close proximity in the complex). Additionally, to account for the possibility of several, spatially distant residues contributing to the complex formation, we disabled the option to automatically define passive residues around the active residues, while the rest of the algorithm options remained default.

An unbiased model for the Arc-PABPN1 complex was made using AlphaFold2 (PMID 34265844) on the ColabFold site (PMID 35637307). As input, the sequences of the human Arc C-lobe and PABPN1 were used to build a model formed of a PABPN1 homodimer bound to Arc C-lobe.

### *In vitro* hippocampal neuronal cell culture

Primary rat hippocampal neurons obtained at postnatal day 0 were cultured in 24-well plates (VWR) on 12 mm glass coverslips (VWR) coated with poly-D-lysine (Sigma Aldrich) and laminin (Roche). Upon tissue homogenization, cells were counted in a Bürker chamber and seeded at the density of ∼75 000 cells per well. After 1.5-2 h incubation plating medium (MEM supplemented with 10 % fetal bovine serum, 0.45 % glucose, 2 mM glutamine and penicillin/streptomycin 100 U/ml–100 µg/ml) it was replaced with defined maintenance medium (neurobasal supplemented with B27, 2 mM glutamine and penicillin/streptomycin 100 U/ml–100µg/ml). Cells were cultured in an incubator with a humid, 5 % CO2 containing atmosphere at 37 ⁰C.

### Silencing of Arc expression

The Arc silencing 29mer shRNA sequence: CTCACGCCTGCTCTTACCAGCGAGTCAGT in retroviral GFP containing vector (Gene ID = 54323) obtained from OriGene (TG7-10356). was delivered to cultured neurons at DIV4 using Lipofectamine3000 (Thermo Fisher Scientific) according to manufacturer instructions. Cells were incubated with Lipofectamine and a transfecting medium (OptiMEM, 2 % B7 Supplement, 1 mM pyruvate, 0.5 mM GlutaMAX, 25 µM β-mercaptoethanol) for 45 min at 37 °C, 5% CO_2_ in a humidified incubator. Next, the medium was replaced with conditioned maintenance medium.

### Chemical LTP

Chemically induced long-term potentiation of synaptic transmission (cLTP) was induced as described (Otmakhov et al., 2004; Szepesi et al., 2013). Hippocampal neuronal cultures (DIV13-15) were exposed to medium containing 50 μM forskolin (Sigma Aldrich), 50 nM rolipram (Sigma Aldrich), and 200 μM picrotoxin (Sigma Aldrich) for 2 hr and immediately fixed with 4 % PFA. All chemicals were dissolved in DMSO and control cultures received DMSO solvent only.

### Immunofluorescence on neuronal cultures

The DIV 13-14 cells growing on coverslips were fixed using 4 % PFA freshly prepared from a 16 % stock (Thermo Fisher Scientific, dissolved 1:4 in 1xPBS) for 10 min RT. Next, PFA was exchanged for 1 x PBS, and the slides were kept at 4 °C until further use. To prevent bacteria growth, sodium azide was added to 0.002 % concentration. Coverslips were next washed in 1xPBS, permeabilized with 0.35 % TritonX100 (Sigma Aldrich) in 1xPBS for 3 min at RT and blocked in 5 % bovine serum albumin (BSA) diluted in 0.1 % TritonX100 in 1xPBS (PBST). Incubation with primary antibodies (list and dilutions in Table 1) diluted in 5 % BSA in PBST was carried for 1 h, at RT. Signals were detected sequentially with secondary antibodies conjugated with fluorophores (list and dilutions in Table 1) diluted in 5 % BSA in PBST. Samples were counterstained with Hoechst 33342 (Invitrogen, H3570) for 10 min RT, washed 3x for 5 min with PBST and mounted on glass slides using Vectashield (Vector Laboratories) and analyzed using a confocal microscope.

### Confocal Microscopy

Fluorescence images were captured using Zeiss LSM800 Airyscan or LSM780 Microscope (Zeiss,) equipped with 63x PlanApo oil immersion objective (NA 1.4). Pixel size was set to 70 nm in xy-direction and 210 nm in z-direction, according to the Nyquist criterion. Confocal pinhole was set to 1 Airy unit for each channel. Fluorescence was excited with the following lasers: 405 nm diode, 488 nm argon ion (LSM 780) or diode (LSM800,) 561 nm DPSS, 640 nm diode (LSM800) or 633 nm HeNe (LSM780). Cross-talk between the fluorophores was eliminated by adjustment of the spectral ranges of the PMT detectors and sequential scanning of the images.

### Quantitative Image Analysis

Channels in the images were aligned using Zen software (Zeiss, Germany), using 0.5 μm fluorescent beads as the reference standard. All quantifications were performed using self-developed PartSeg Software (Bokota et al., 2021) in 3D images. Neuronal nuclei were distinguished based on NeuN neuronal marker staining and segmented based on DNA staining. PABPN1 staining was segmented by multi-thresholding using the Multiple Otsu algorithm (Liao et al., 2001), which separated the pixels of a 3D image into 7 classes according to the increasing gray levels intensity of the staining within the image. Class 1 covered pixels with the minimal intensity values, class 7 represented pixels with the highest pixel intensity values. The volume of nuclear speckles (combined classes 3-7, which corresponded to specific staining with SF3a66-speckle marker, not shown) was expressed as a percentage of nucleus volume.

## Results

### Localization of neuronal activity induced Arc to the interchromatin space of dentate granule cell nuclei in vivo

We first examined nuclear localization of Arc following seizures (Fig. 1A) and LTP induction in the DG in vivo (Fig. 1B). Immunofluorescent staining of hippocampal sections from untreated control rats showed overall paucity of nuclear Arc expression with clear fluorescence observed in sparsely distributed hippocampal pyramidal cells and dentate granule cells, consistent with the sparse all-or-none pattern of Arc mRNA expression (Lyford et al., 1995). A 50-fold increase in average Arc nuclear signal in granule cells was observed by 2 h after PTZ (Fig. 1C and 1E). Unilateral HFS (400 Hz, 8-pulse burst pattern) of the medial perforant input to dentate gyrus induced stable LTP of synaptic transmission (Fig. 1B), and enhanced Arc expression in granule cell nuclei (Fig. 1D and 1F). Nearly all granule cells in dorsal DG exhibited strong Arc immunofluorescence, with mean increases in signal intensity over single nuclei of 55-fold and 40-fold at 1 h and 3 h, respectively, relative to the contralateral non-stimulated DG (Fig. 1F). Arc expression was also observed at lower signal intensity in the perinuclear cytoplasm. As previously shown, Arc was selectively expressed in granule cells with no detectable expression in other cell types of the dentate gyrus (interneurons, mossy cells, glia). Notably, the nuclear Arc immunofluorescence was punctate and non-overlapping with DAPI stained chromatin, indicating primary localization of activity-induced Arc to the interchromatin space (Fig. 1C and 1D).

**Figure 1.**
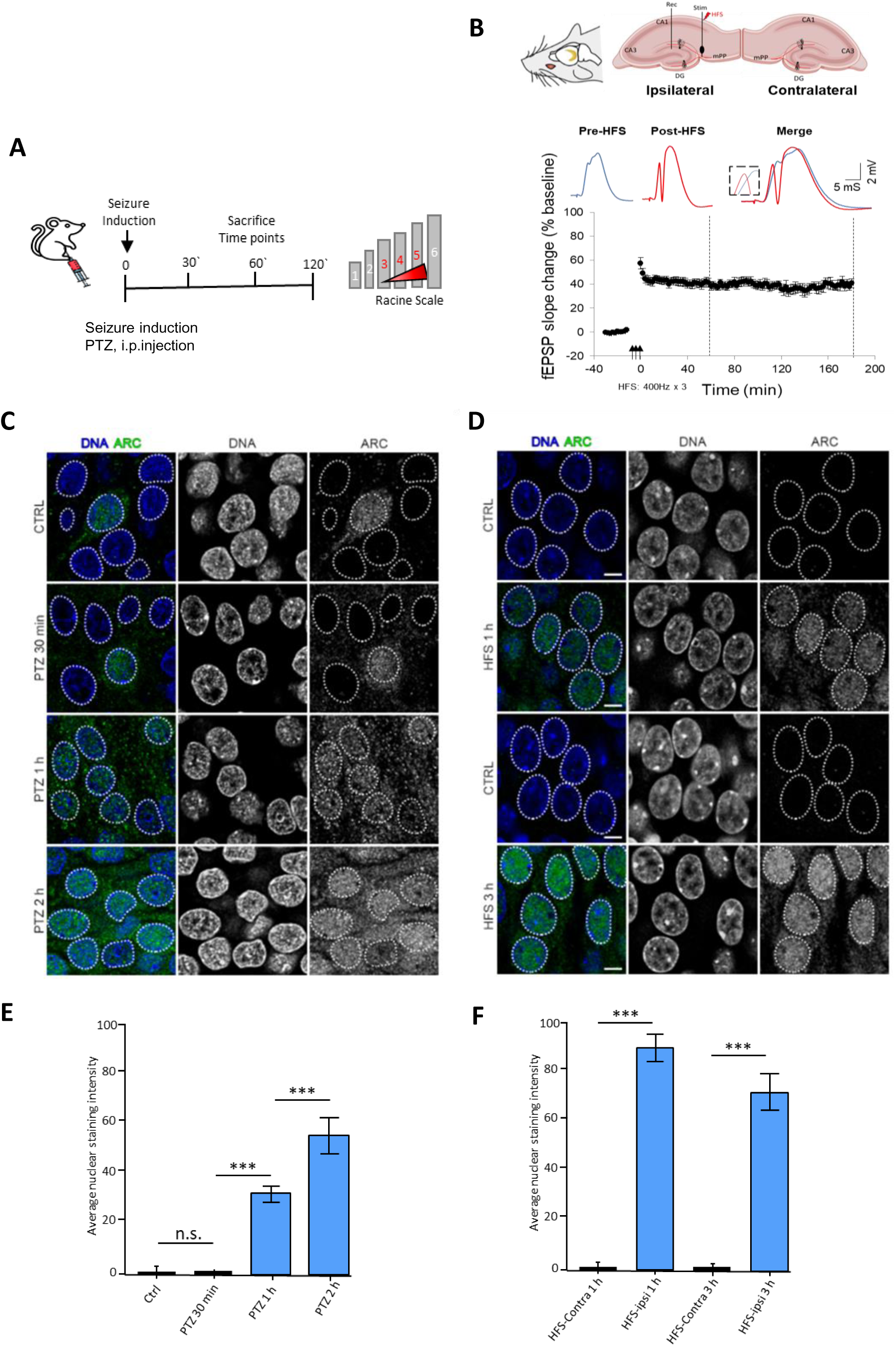
Localization of neuronal activity induced Arc to the interchromatin space of dentate granule cell nuclei in vivo. (A) The modified Racine scale for the behavioral scoring of seizure severity in rats following a single i.p. injection of PTZ. The grade scale is applied for seizure-related behavioral changes and assessment of seizure severity (see Materials and Methods section for details). Each number represents a different degree of seizure severity. Rats were killed at 30 min, 1 h, and 2 h after PTZ injection leading to generalized seizures (grades 3-5). (B) Top panel. Schematic illustration of the electrode placement for unilateral induction of LTP at the medial perforant path input to dentate granule cells in vivo. Middle panel. Representative field potentials from baseline period and the last 5 minutes of post-HFS recording. Bottom panel. Time course plots of fEPSP slope values recording before and after HFS (indicated by arrows). Values mean ± SEM in percent of baseline. Rats were killed at 1 h and 3 h post-HFS (stippled line). (C) Coronal sections (focal plane, 210 nm) through DG neurons at 30, 60 and 120 minutes after PTZ injection. The outline of the nuclei borders is marked with a dashed line. Single plane from confocal images with the Arc immunostaining (depicted in green and gray on single channel pictures), and chromatin staining (depicted in blue and gray on single channel pictures) are shown. Bars represent 5 μm. (D) Coronal section through granule cell region from ipsilateral, HFS-treated DG and contralateral DG control. Color labeling as in panel C. Bar scale 5 μm. (E) Quantitative analyses of Arc signal accumulation in the nuclei of granule cells following PTZ induced seizure activity. Bar graphs show average Arc immunofluorescence intensity ± SEM (n=3, n=10), *** p-value < 0.001. Non-parametric two-tailed, unpaired t-test. (F) Quantitative analyses of Arc signal accumulation in the nuclei of granule cells at 1 h and 3 h post-HFS. Graphs show changes in nuclear Arc immunofluorescence signal intensity as mean ± SEM (n=10), stars represent p-values < 0.001 *** of the non-parametric two-tailed, unpaired t-test.

### Enhanced Arc expression in the DG nucleosol subcellular fraction during LTP

Subcellular fractionation of DG tissue into cytosol, membrane, cytoskeletal, nuclear soluble (nucleosol) as well as nuclear chromatin-bound fractions was used to investigate Arc localization in non-stimulated tissue and after LTP induction (Fig. 2A). Immunoblotting with nuclear compartment marker proteins confirmed enrichment of matrin 3 in the nucleosol and histone 1 in the chromatin-bound fraction (Fig. 2B). In non-stimulated DG, Arc is enriched 2-3-fold in the cytoskeletal fraction relative to cytosol and membrane fractions (Fig. 2B and 2C). Within the nucleus, Arc immunoreactivity was consistently detected as a low-intensity band in the nucleosol, whereas Arc was only scarcely detectable in the chromatin-bound fraction. Following LTP induction, at 2 h post-HFS, Arc expression was significantly enhanced in all fractions from the ipsilateral DG relative to contralateral control (Fig. 2B and 2C), with the largest mean increase of 6-fold detected in the nucleosol. Although Arc immunoreactivity was also significantly increased in the chromatin-bound fraction post-HFS, Arc expression remained very low. Taken together the results show massive induction and accumulation of Arc in the interchromatin space of postsynaptic granule cells and the DG nucleosol fraction.

**Figure 2.**
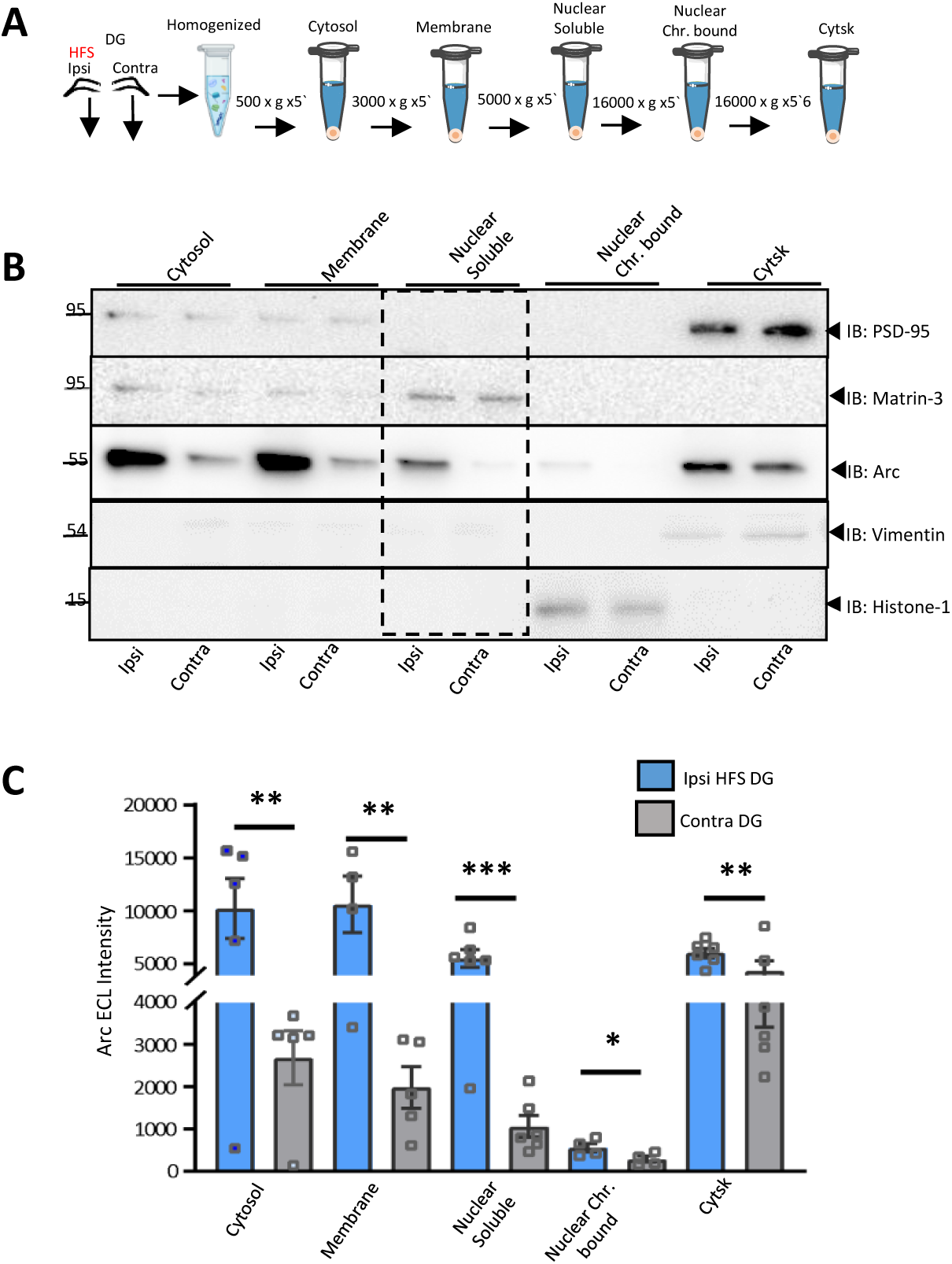
Enhanced Arc expression in nucleosol fraction during DG LTP in vivo. (A) Schematic representation of the subcellular fractionation protocol of DG tissue from the rat brain 2 hours post-HFS *in vivo*. Cytosol, membrane, nuclear soluble (nucleosol), nuclear chromatin bound, and cytoskeletal fractions were obtained from HFS-treated, ipsilateral (Ipsi) DG and contralateral (Contra) non-stimulated DG. (B) Immunoblot analysis of subcellular fractions from HFS-treated, ipsilateral (Ipsi) and contralateral (Contra) DG. Fractions were validated using compartment-enriched marker proteins: PSD-95, postsynaptic cytoskeletal fraction; Matrin-3, nucleosol fraction; Vimentin, cytoskeletal fraction; Histone-3, nuclear chromatin fraction. 40 µg protein loaded per lane. *n=5*. (C) Bar graphs show densitometric quantification ECL intensity from Arc immunoblots. Values are mean (± SEM) Significant difference between HFS-treated and contralateral DG are indicated *p<0.05; **p<001. ***p<0.0001, Mann-Whitney *t*-test. n=5

### Mass spectrometric proteomic analysis of nuclear Arc complexes identifies proteins with roles in mRNA 3’-end processing and polyadenylation

To screen for binding partners, Arc was immunoprecipitated from the nucleosol fraction of ipsilateral HFS-treated and contralateral DG and subjected to mass spectrometric proteomic analysis. We first confirmed by immunoblot analysis enhanced Arc expression in the nucleosol input sample from HFS-treated DG relative to control. Immunoprecipitation enriched for Arc, which exhibited a 5-fold increase in abundance post-HFS (Fig. 3A and 3B). MS Proteomic analysis was performed on samples from a separate group of rats. Arc immunoprecipitated was gel-extracted, trypsin digested, and target proteins were identified by LC-MS/MS. We identified unique proteins with a protein False Discovery Rate of 1% (Supplemental Table 1, PRIDE dataset identifier PXD053124). STRING database analysis for known and predicted protein-protein interactions identified 3′-end mRNA processing and polyadenylation regulation as key categories for Arc interactors (Fig. 3C).

**Figure 3.**
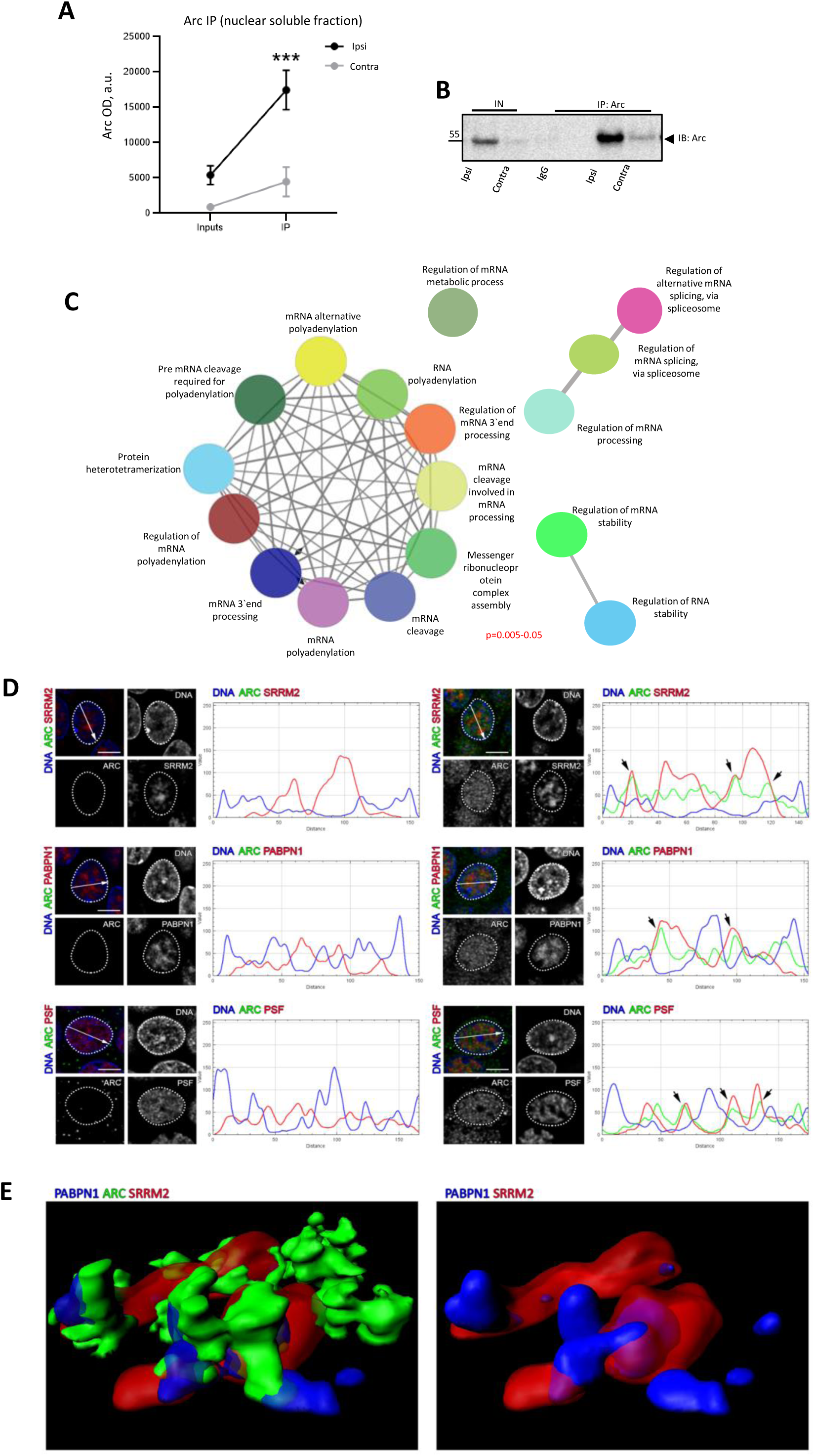
Mass spectrometric proteomic analysis of immunoprecipitated nuclear Arc complexes identifies proteins with roles in mRNA 3’-end processing and polyadenylation. Arc immunoprecipitated complexes from DG nucleosol (100 ug) were separated by SDS-PAGE. (A) Area chart showing enhanced Arc protein during HFS-LTP induced in rat dentate gyrus comparing ipsilateral compared to contralateral. *n* = 8; Student’s *t*-test, \**p* < 0.05. (B) Representative immunoblot of Arc in nucleosol input and Arc immunoprecipitate from HFS-treated and contralateral DG. (C) The results obtained from mass spectrometry analysis showing higher levels of pre-mRNA proteins. (D) Neuronal nuclei from the dentate gyrus of hippocampal formation of a control rat (left side panels) and animals subjected to PTZ-induced seizures (right side panels) 2 hours post seizure induction. Single plain confocal images show immunostainings for SRRM2 (top panels), PSF (middle panels) and PABPN1 (bottom panels) in red, Arc protein in green and counterstaining of DNA in blue on the merged pictures and RGB profiles. RGB profiles show signal intensity values (0-255 range in the y-axis) across white arrows depicted on the merged images. Black arrows show signal overlap between Arc and depicted protein. Bars represent 5μm. (E) Reconstruction of nuclear speckle based on 3D image of immunofluorescent signal. Neuronal nucleus was imaged using confocal microscope in the dentate gyrus of hippocampal formation of an animal subjected to PTZ-induced seizures. Volumes occupied by SRRM2 (nuclear speckle marker), PABPN1 and ARC protein are represented in red, blue, and green colors respectively.

For validation of Arc interaction partners, we selected 3 proteins with roles in mRNA polyadenylation and processing. 1) PolyA-binding protein nuclear 1 (PABPN1; also known as polyA-binding protein 2), is a multifunctional protein with primary roles in pre-mRNA 3’-end polyadenylation and splicing (Muniz et al., 2015). 2) Serine/Arginine Repetitive Matrix Protein 2 (SRRM2) is a major component of nuclear speckles, which are membrane-less nuclear bodies enriched in splicing factors. SRRM2 is linked to nuclear speckle formation and organization of splicing condensates (Ilık et al., 2020; Faber et al., 2022; Ilık and Aktaş, 2022; Xu et al., 2022), 3) Polypyrimidine splicing factor (PSF), also known as splicing factor protein and glutamine rich (SFPQ), is a component of paraspeckles, with ascribed roles in splicing, DNA damage repair, and transcriptional regulation (Lee et al., 2015; Lim et al., 2020).

### Seizure-induced Arc coincides with PABPN1, PSF, and SRRM2 in granule cell nuclei

We evaluated subnuclear localization of Arc by immunofluorescence and confocal analysis of dentate granule cell nuclei following PTZ-evoked seizure activity. Arc immunofluorescence was detected in nuclei mostly after neuronal activation by PTZ, where it coincided with nuclear speckle markers PABPN1 and SRRM2 (Fig. 3D and 3E), and PSF protein (Fig. 3D). Arc immunofluorescence signal colocalized with PABPN1 and SRRM2 either at the border of large nuclear speckles, or within smaller nuclear speckles (Fig. 3D; depicted by arrows on RGB profiles where Arc is shown in green, and PABPN1 and SRRM2 are shown in red). In control animals, PSF formed discrete and numerous nuclear domains more homogeneously distributed within the nucleus. Upon induction of seizures, PSF formed larger foci, most of which coincided with Arc immunofluorescence signal (Fig. 3C; depicted by arrows on RGB profiles where Arc is shown in green, and PSF is shown in red). Figure 3E shows a 3D reconstruction of Arc localization to a nuclear speckle, where PABPN1, SRRM2 and Arc are shown in blue, red and green respectively.

### Arc interacts with PABPN1 and PSF in the nucleosol and exhibits enhanced complex formation during in vivo LTP

The subcellular distribution and impact of LTP induction on the expression of PABPN1, PSF, and SRRM2 in the adult brain is unknown. Therefore, we first examined the expression of the putative binding partners in DG subcellular fractionations by immunoblot analysis (Fig. S1). In control unstimulated DG tissue, PABPN1 is abundant in the nucleosol and chromatin-bound fractions but absent from remaining fractions. PSF was abundant in the nucleosol but absent from the chromatin fraction, with low levels of expression in the cytosol and membrane fractions, consistent with PSF as a nuclear paraspeckle protein and constituent of dendritic RNA transport granules (Kanai et al., 2004). SRRM2 was also predominantly located in the nucleosol, with only a weak band detected in the chromatin fraction. LTP induction did not result in qualitative changes in the subcellular distribution of the partner proteins as measured 1 h post-HFS (Fig. S1). However, within the nucleosol compartment, PABPN1 exhibited a significant 1.5-fold increase in expression along with a 5-fold elevation in Arc, whereas SRRM2 and PSF levels were not different from control (Fig. 4A). The results indicate that LTP-inducing synaptic stimulation does not alter the compartmental distribution of PABPN1, PSF, and SRRM2 but enhances expression of PABPN1 in the nucleosol.

**Figure 4.**
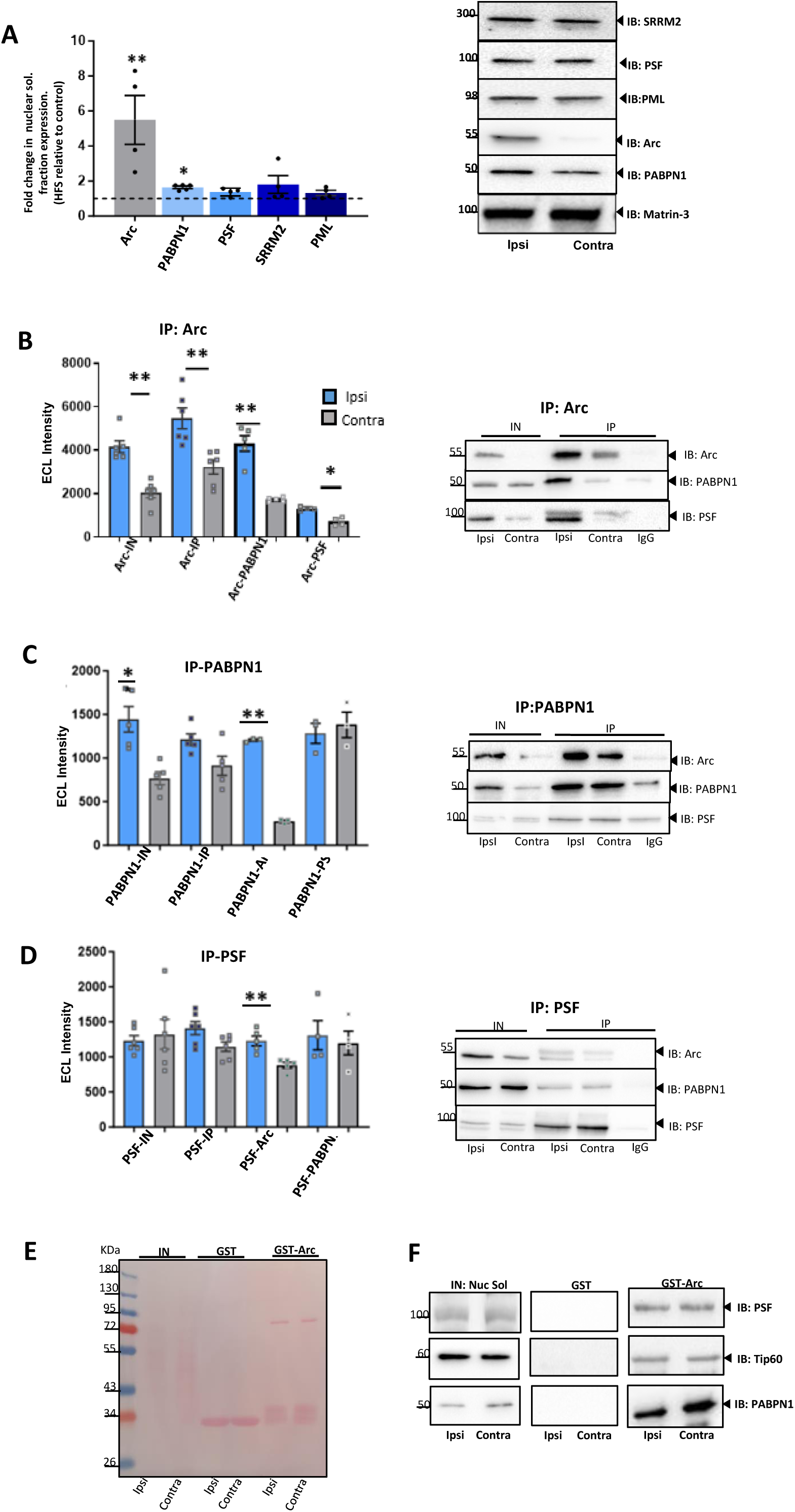
Arc interacts with PABPN1 and PSF in the nucleus during DG LTP in vivo. (A) Densitometric quantification of immunoblots showing fold change in protein expression in the nucleosol fraction of ipsilateral HFS-treated DG relative to the contralateral DG. \**p* < 0.05, \*\**p* < 0.001, Mann-Whitney *t*-test; *n* = 5. (B-D) Co-immunoprecipitation analysis. Bar graphs show quantification of immunoblot signal from DG nucleosol input sample and immunoprecipitated from HFS-treated and contralateral DG. *Significant difference between ipsilateral HFS-treated and contralateral DG. *n* = 5; Mann-Whitney *t*-test, *p < 0.05, **p < 0.001. (B) Arc immunoprecipitation. (C) PABPN1 immunoprecipitation. (D) PSF immunoprecipitation. (E) DG nucleosol fraction was incubated with GST-fused Arc or GST alone bound to glutathione-Sepharose beads. Ponceau protein staining. (F) Immunoblots for PSF, TIP60, and PABPN1 in nucleosol input, GST control, and GST-Arc pulldown.

To probe Arc protein-protein interactions, we performed co-immunoprecipitation analysis in the nucleosol fraction from HFS-treated and contralateral DG tissue. In Arc immunoprecipitated samples (Fig. 4B), a PABPN1 immunoreactive band was detected at 50 kDa in non-stimulated DG, which exhibited a 2.3-fold increase in intensity in HFS-treated DG. PSF also co-immunoprecipitated with Arc in unstimulated DG and showed a significant 1.5-fold increase post-HFS (Fig 4B). A faint band in the IgG-bead control lane also indicates weak non-specific binding of PSF to beads, whereas PSF levels in Arc immunoprecipitate were consistently and significantly higher (Fig. 4B). We also examined co-immunoprecipitation of SRRM2 with Arc but found no consistent evidence for complex formation using a variety of anti-SRRM2 antibodies and buffer conditions. We therefore focused the reciprocal co-immunoprecipitation analysis on PABPN1 and PSF. In PABPN1 immunoprecipitated samples (Fig. 4C), Arc was detected in control DG and exhibited a 7-fold increase in HFS-treated DG. PSF also co-immunoprecipitated with PABPN1, but there was no difference between control and HFS-treatment. In PSF immunoprecipitated samples (Fig. 4D), Arc and PABPN1 were both detected in control and HFS-treated DG, but only Arc showed significantly enhanced co-immunoprecipitation with PSF post-HFS. RNase A treatment of DG tissue samples did not inhibit co-immunoprecipitation of PABPN1 or PSF with Arc, demonstrating capture of protein-protein interaction complexes rather than indirect association mediated by RNA binding (Fig. S2). We then performed pulldown experiments in the nucleosol fraction using human recombinant GST-Arc as bait (Fig. 4E and 4F). Clear immunoblot signals for PABPN1 and PSF were obtained in the GST-Arc pulldown, with particularly robust recovery of PABPN1 (Fig. 4F). As a positive control, we also detected a previously identified Arc interaction partner, the histone acetyl transferase TIP60 (Wee et al., 2014) (Fig. 4F). None of the partner proteins were detected when samples were incubated with glutathione-coupled Sepharose beads as a control for non-specific binding (Fig. 4F).

**Figure 5.**
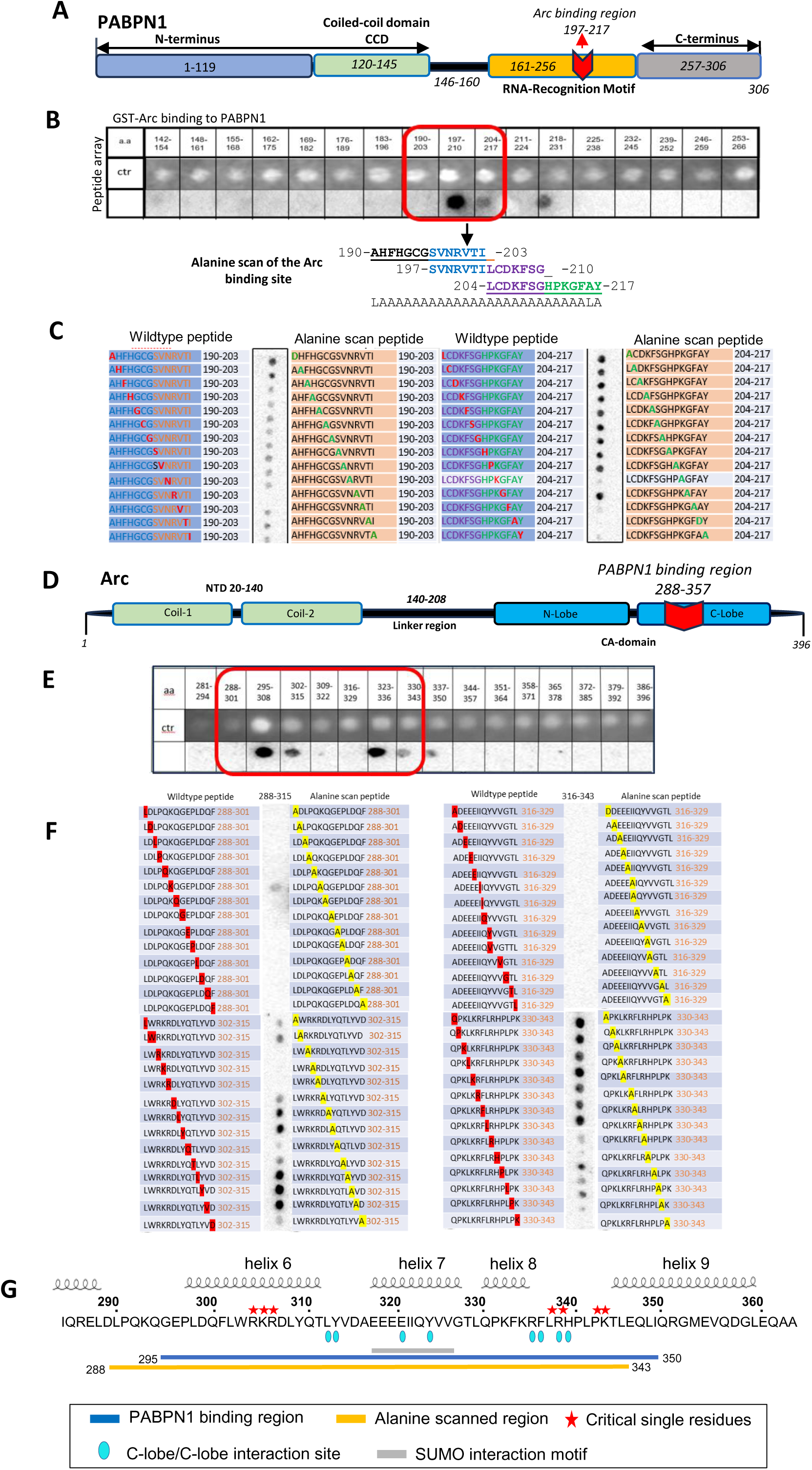
Arc binds to PABPN1 polyA RNA recognition motif. Peptide biding array analysis Arc interaction with PABPN1. Cellulose-bound peptide arrays covering the human PABPNI or rat Arc sequences were incubated with purified GST-fused Arc or GST-PABPN1, respectively. (A) Cartoon of PABPN1 domain structure indicating position of Arc binding site. (B) Sample peptide array of GST-Arc binding to PABPN1 peptides. The red box indicates the Arc binding region (197-217) used for alanine scanning. (C) Alanine scanning of GST-Arc binding region on PABPN1. Wildtype and alanine scan peptides shown. (D) Cartoon of Arc domain structure indicating putative PABPN1 binding site. (E) Sample peptide array of GST-PABPN1 binding to Arc peptides. The red box indicates the Arc C-lobe binding region (288-357) used for alanine scanning. (F) Alanine scanning of GST-PABPN1 binding region on Arc C-lobe. Wildtype and alanine scan peptides are shown. (G) Amino sequence of C-lobe region with position of helices 6, 7, 8, and 9. The PAPBN1 binding region and position of critical single residues are indicated. The region of PABPN1 binding contains several conserved dimerization sites of C-lobe/C-lobe interact and putative SUMO interaction motif.

In sum, co-immunoprecipitation and GST-Arc pulldown analysis demonstrated that Arc interacts with PABPN1 and PSF in the nucleosol compartment. The interaction is detected under basal conditions, with enhanced complex formation following LTP induction and accumulation of Arc in the nucleus. The reciprocal immunoprecipitation between PABPN1 and PSF also suggests that Arc, PABPN1 and PSF may occur together in the same complex. Next, we sought to identify a possible direct binding of Arc to a nuclear partner. We considered PABPN1 as the best candidate direct binder, given its robust co-purification with Arc and recovery in the GST-Arc pulldown.

### Interaction between the Arc capsid domain and PABPN1 RNA recognition motif

Direct binding of Arc to PABPN1 was examined using peptide binding arrays. Purified human GST-Arc prepared in its monomeric state was used to probe a peptide tiling array covering the 306 amino acid full-length sequence of rat PABPN1 (Fig. 5). PABPN1 has multiple roles in 3’-end RNA processing and metabolism, including regulation of polyadenylation, poly(A) site selection, RNA degradation and RNA export (Banerjee et al., 2013). PABPN1 has three characterized domains: an RNA recognition motif (RRM) that binds poly(A) RNA, a coiled coil domain (CCD) which stimulates polyA polymerase activity during polyadenylation, and a C-terminal domain which together with the RRM facilitates polyA binding and PABPN1 oligomerization (Fig. 5A). We found that Arc binds with high affinity to a single 14 amino acid stretch, from residues 197 to 210, and with lower affinity to the overlapping 207-217 peptide and 218-231 peptide, in the middle of the PABPN1 RRM (Fig. 5B and Fig. S3). To identify critical residues for binding we performed alanine scanning with single alanine substitutions along the 190-217 segment of the RRM. Arc binding was reduced by point mutations from 192 to 203 and essentially abolished by R200A, K213A, F215A, A216D, and Y217A substitutions (Fig. 5C).

We similarly sought to identify PABPN1 binding sites on Arc. Mammalian Arc has two major domains separated by an unstructured linker region (Fig. 5D). The NTD is a predicted antiparallel coiled coil involved in Arc membrane binding, endocytosis, and oligomerization including capsid formation (Hallin et al., 2018; Eriksen et al., 2021). The CTD, or capsid (CA) domain, has extensive structural homology with the capsid domain of retroviral Gag polyprotein (Zhang et al., 2015). Arc CA is comprised of two lobes, the N-lobe and C-lobe. The N-lobe harbors a hydrophobic pocket that mediates Arc binding to several postsynaptic partner proteins (Zhang et al., 2015; Nielsen et al., 2019; Hallin et al., 2021). Both lobes have conserved dimerization motifs implicated in higher-order Arc oligomerization (Zhang et al., 2019). Purified PABPN1 bound with high affinity to peptides of the Arc NTD and CA, whereas no binding was detected to peptides from the unstructured linker region or termini (Fig. 5E and Fig. S3). In the N-terminal coiled coil, strongest binding was observed in overlapping peptides from aa 28-56 in the middle of Coil-1. Within the CA domain, binding was observed for a single peptide (aa 211-224) in the N-lobe ligand binding region and for multiple overlapping peptides in the 295-350 aa segment spanning helices 6, 7, and 8 and of the C-lobe (Fig. 5E and Fig. S3). For alanine scanning analysis we focused on the prominent PABPN1 binding to the C-lobe (Fig. 5F). Single substitutions that abolished PABPN1 binding to peptide 302-315 (R304, K305, R306, Q310) and peptide 330-343 (L337, R338, P342, K343) were identified (Fig. 5F and 5G). Wildtype peptides that were negative for PABPN1 binding (288-301 and 316-329) remained negative in the mutant peptides. The critical single residues occurred in patches of 2 or 3 consecutive residues in helix 6 and along the segment connecting helix 8 and 9 (Fig. 5F and 5G). Notably, the region of the C-lobe that binds PABPN1 harbors eight predicted dimerization sites for C-lobe/C-lobe intersubunit binding (Fig. 5G). The segment connecting helix 8 and 9 has four dimerization sites, and one of these sites (R338), is also a critical single residue for PABPN1 binding (Fig. 5G).

### Structural modeling of Arc C-lobe interaction with PABPN1 RRM

To gain insight into the structural basis of Arc-PABPN1 recognition, we used the HADDOCK2 web server to perform protein-protein docking analyses (van Zundert et al., 2016). Briefly (see Methods), we used a stretch of residues (identified as contributing to complex formation by peptide array analysis; Figure 6A, 6B, and Fig. S4) as experimental constraints in a panel of HADDOCK docking runs. Anchor residues (i.e., those predicted to be in close spatial proximity in the complex) were defined as non-overlapping 3-residue stretches in the 302-358 residue region for Arc and the Lys213 residue for PABPN1, respectively. The resulting HADDOCK energy estimates showed significant energy differences between runs initiated with different residue ranges as anchor points (Figure 6C). Further visual inspection of the modeled complexes revealed that the top 3 energy values (Figure 6C, highlighted in red) correspond to nearly identical structures (within 1.4Å RMSD), indicating convergence of the docking algorithm to the unique solution (Figure 6D). Structural analysis of the modeled complexes indicates that a negatively charged loop in the C-lobe (aa D315-Q323) promotes complex formation by providing an extensive hydrogen bonding network with residues K207, S209, and K213 of PABPN1 (Figure 6D and 6E).

**Figure. 6.**
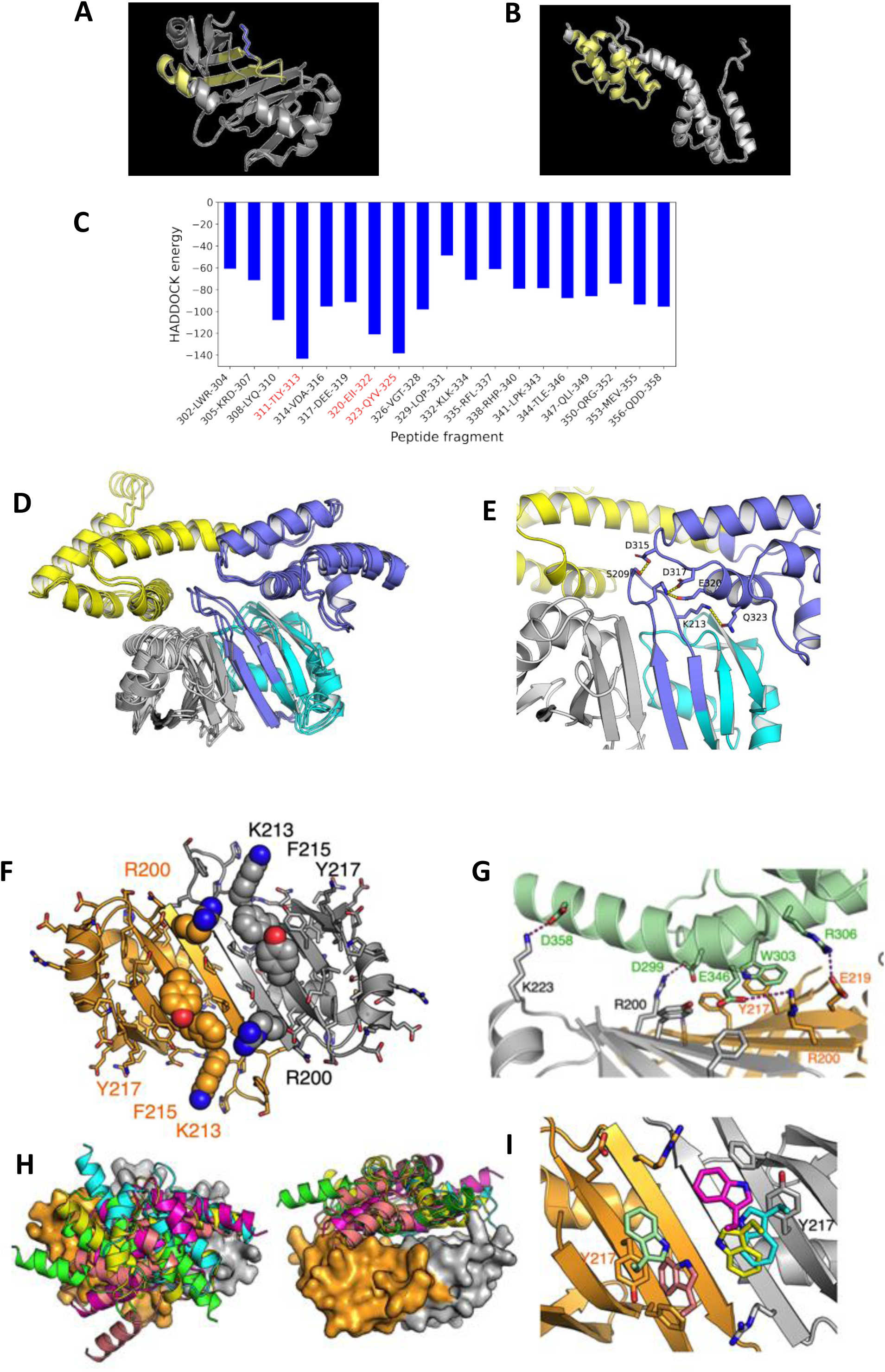
Structural models of Arc C-lobe binding to the PABPN1 RRM. A) Human PABPN1 structure (PDB: 3B4M) used in the HADDOCK runs. Residues involved in complex formation (according to the peptide array results) are highlighted in yellow, while the most prominent residue, Lys213, as indicated by the alanine scanning results, is highlighted in blue. B) Rat Arc structure (PDB: 6GSE) used in the HADDOCK runs. Residues involved in complex formation (according to peptide array results) are highlighted in yellow, (C) Distribution of HADDOCK energy estimates for the complexes obtained by docking runs initiated with different sets of residues defined as interacting residues. The top 3 energy values are highlighted in red and the corresponding complex structures are visualized in panel D. (D) Ensemble of the top 3 scored HADDOCK complexes. Arc shown in yellow. Monomers of PABPN1 are shown in blue (interacting monomer) and gray (non-interacting monomer). Regions identified as involved in the interactions by the peptide array experiment (288-360 and 197-217 for Arc and PABPN1, respectively) are shown in purple. (E) Structural insights into the interactions that maintain the modeled ARC-PABPN1 complex. The coloring follows the scheme of panel D. Interacting residues are shown as sticks. (F) Predicted models for the Arc C-lobe/PABPN1 complex from AlphaFold2. The PABPN1 dimer, viewed along the symmetry axis. Key residues identified in alanine scanning are labelled and shown as spheres. The two protomers are colored orange and grey. (G) Salt bridge and aromatic interactions at the interface for the top ranked model. In addition, several hydrogen bonds and van der Waals interactions are present (not shown). Arc C-lobe is shown in green. (H) Superposition of all five AlphaFold2 models for the complex indicates that Arc C-lobe (cartoons) is predicted to always bind the surface shown in panel F. Left: top view (same as in F), right: side view. (I) In all predicted complexes, Trp303 from Arc C-lobe is central to the binding interface, always directly interacting with Tyr217 of PABPN1.

An unbiased model for the Arc-PABPN1 complex was made using AlphaFold2 based on amino acid sequence alone and without any contact restraints. Very similar results were obtained using AlphaFold3, which recently became available (Abramson et al., 2024). The top five models for the Arc C-lobe/PAPBN1 complex all show the Arc C-lobe bound onto the same surface of the RRM in the PABPN1 homodimer (Fig. 6F). Key residues identified in alanine scanning form a large part of this interface, including PABPN1 R200, K213, F215, and Y217. As a result, the predicted binding surface has several options for π-interactions in the middle and a positively charged rim. These properties result in a model with a central aromatic stacking interaction and several salt bridges, as exemplified for the top ranked model (Fig. 6G). Superposing all five models indicates that Arc C-lobe consistently is predicted to binds onto the same surface, with minor variation in orientations (Fig. 6H). W303 of the Arc C-lobe is always in the middle of the binding interface, directly interacting with Y217 from one of the PABPN1 protomers (Fig. 6I). W303 in helix 6 is adjacent to a patch of residues (R304, K305, R306) critical for binding based on the peptide analysis, with the AlphaFold2 model also showing R306 interaction (Fig. 6G). Interaction surfaces on helices 8 and 9 are also consistent with the peptide array data. Taken together, the Haddock and AlphaFold models corroborate the experimental peptide array results on C-lobe binding to the PABPN1 RRM at the level of identified critical residues but suggest different possible binding modes. Deciphering the exact binding mode will require experimental structure determination.

### Arc regulates basal and neuronal activity-dependent formation of PABPN1 nuclear speckles

Synaptic plasticity requires de novo gene expression and post-transcriptional nuclear processing of mRNA. PABPN1, a component of nuclear speckles, is implicated in many aspects of post-transcriptional regulation in addition to its canonical role in polyadenylation regulation. Nuclear speckles are dynamic nuclear bodies, which undergo relocalization, fusion, and morphological changes in response to cell stimuli (Grabowska et al., 2022; Szczepankiewicz, 2024). We considered that Arc, as a direct binding partner of PABPN1, could play a role in neuronal activity-dependent reorganization of nuclear speckles.

We, therefore, examined PABPN1 and Arc by immunofluorescent staining following chemical LTP (cLTP) treatment of primary hippocampal neuronal cultures (Fig. 7). cLTP treatment enhanced the expression of Arc and c-FOS, another immediate early gene product, in neuronal nuclei (Fig. 7A). PABPN1 occurred predominantly in the interchromatin space (depicted by arrows on RGB profile in Fig. 7B) where it coincided with Arc fluorescent signal. Bright PABPN1 foci colocalized with the nuclear speckle marker SF3a66 (Fig. 7C) and were surrounded by a weaker and homogeneous signal dispersed in the nucleoplasm (Fig. 7C) depicted with green arrows on RGB profiles). To investigate a possible role for Arc in regulation of nuclear speckles, we depleted Arc protein by a specific shRNA (Fig. 7D). Using 3D image segmentation (Fig. 7E), we quantified the number, surface area and volume of nuclear speckles based on PABPN1 immunofluorescence (Fig. 7E). In control, non-transfected cells from the same cultures, cLTP treatment induced a significant increase in the number, total surface area, and total volume of nuclear speckles, with no change in their mean volume (Fig. 7F). Further analysis of speckles by size category showed increases in the number of small (<2 µm^3^) and very large (>18 µm^3^) speckles upon stimulation (Fig. 7G). Depletion of Arc under basal conditions resulted in more numerous, slightly smaller speckles. cLTP treatment did not further increase speckle number in Arc depleted neurons, but significantly increased speckle volume and surface area. Interestingly, depletion of Arc diminished cLTP-associated formation of small (<2 µm^3^) nuclear speckles. All the above indicates a role for Arc in neuronal activity-dependent formation and reorganization of nuclear speckles.

**Figure 7.**
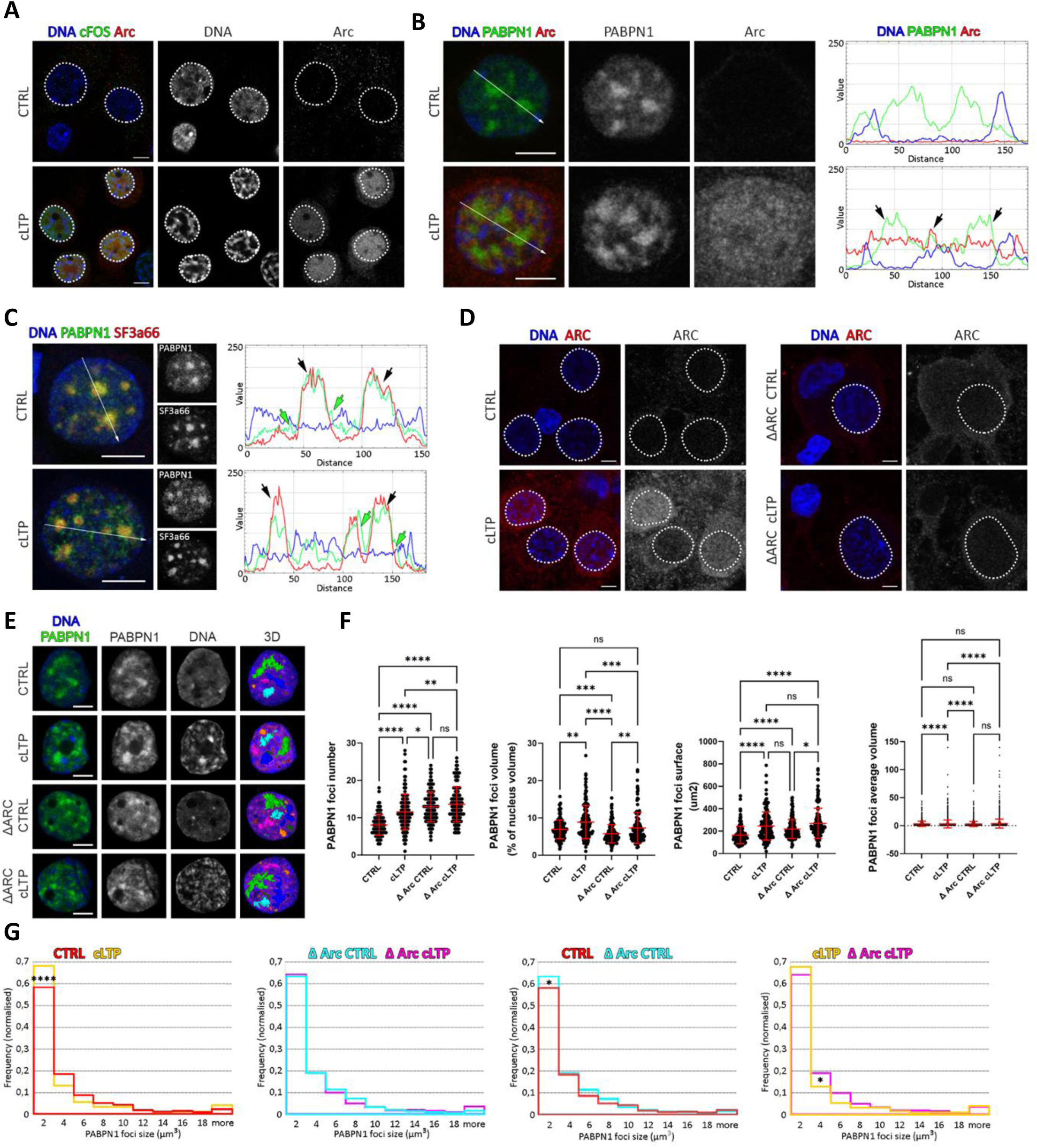
Arc regulates formation of PABPN1 nuclear speckles upon neuronal stimulation. Hippocampal cultures subjected for 2 h to the chemically induced LTP (cLTP) or treated with DMSO vehicle (CTRL B). The bars shown in images represent 5 μm. (A) Middle plain from confocal images shows DNA (depicted in blue on the merged image, and gray on single channel pictures), expression of Arc (depicted in red on the merged image, and gray on single channel pictures), and c-FOS neuronal activation marker (shown in green on the merged image). The outlines of the nuclei borders are marked with a dashed line. (B) Middle plain from confocal images shows DNA (depicted in blue on the merged image), expression of Arc (depicted in red on the merged image, and gray on single channel pictures), and PABPN1 protein (shown in green on the merged image, and gray on single channel pictures). RGB profiles show signal intensity values (0-255 range in the y-axis) across white arrows depicted in the merged images. Black arrows show signal overlap between Arc and PABPN1 protein. (C) Single plain from confocal images shows immunostaining of nuclear speckles marker SF3a66 (depicted in red on the merged image, and gray on single channel pictures), PABPN1 protein (depicted in green on the merged image, and gray on single channel pictures) and counterstaining of DNA (depicted in blue on the merged image). RGB profiles show signal intensity values (0-255 range in the y-axis) across white arrows depicted in the merged images. Black arrows show signal overlap between SF3a66 and PABPN1 protein. Green arrows show PABPN1 protein outside of nuclear speckles. (D) Neuronal cultures transfected with shRNA specific for rat Arc were subjected for 2 h to the chemically induced LTP (cLTP) or treated with DMSO vehicle (CTRL). Cells with silenced Arc expression (ΔArc) were selected based on GFP signal (not shown). Single plains from confocal images show DNA (depicted in blue on the merged image) and expression of Arc (depicted in red on the merged image, and gray on single channel pictures). The outlines of the nuclei borders are marked with a dashed line. (E) Examples of neuronal nuclei of cells from experimental conditions described in D. Single plains from confocal images show DNA (depicted in blue on the merged images, and gray on single channel pictures) and PABPN1 (depicted in green on the merged image, and gray on single channel pictures). The 3D images show segmentation based on strong PABPN1 staining. (F) The quantitative analyses of PABPN1 foci show results from 3 independent experiments, stars represent p-values < 0.0001 ***, < 0.001 ***, < 0.005 **and < 0.01 * of the two-tailed, unpaired Kruskal-Wallis test. (G) Frequency distribution of PABPN1 foci volume shown in E, stars represent p-values < 0.0001 ***, and < 0.01 * of the two-tailed, unpaired Kruskal-Wallis test.

## Discussion

This study identifies PABPN1 as an Arc binding partner in the nucleus and demonstrates enhanced native complex formation during synaptic plasticity in the adult brain in vivo. We further provide evidence for direct binding of Arc to the RRM of PABPN1, and Arc-dependent regulation of basal and neuronal activity-dependent formation of PABPN1 nuclear speckles.

Following LTP-inducing stimulation of the medial perforant path, newly synthesized Arc protein localizes to the granule cell interchromatin space and the nucleosol subcellular fraction, where it forms a complex enriched in proteins involved in RNA processing and metabolism. Reciprocal co-immunoprecipitation analysis demonstrates native Arc interaction with PABPN1 and PSF in the nucleosol. Affinity purification with recombinant GST-Arc further confirms the interaction and supports direct binding between partners. This Arc nuclear complex is detected in naive, unstimulated DG and greatly enhanced following LTP induction. Of the two partners studied, PABPN1 exhibited the strongest biochemical interaction with Arc and LTP-associated regulation. Results from in vitro peptide binding arrays combined with alanine scanning analysis show that recombinant monomeric Arc binds selectivity to the PABPN1 RRM. Given the direct binding to PABPN1 and enrichment of Arc complexes with RNA binding proteins and splicing factors, we explored possible roles for Arc in regulating nuclear speckles. Using immunofluorescence and 3D image analysis in cLTP-treated hippocampal neurons, we show discrete PAPBN1 foci that correspond to SF3a66-labelled nuclear speckles, as well as more homogenous nuclear expression outside of speckles. Similar activity-dependent regulation of speckles is observed following kainic acid induce seizures using various speckle markers (Szczepankiewicz, 2024). Here, we show that neuronal activity-induced Arc occupies the same interchromatin space as PABPN1 but is not concentrated to speckles. cLTP treatment induces reorganization of PABPN1 foci, characterized by increases in the number of small (<2 µm^3^) and very large (>18 µm^3^) foci. Knockdown of Arc increases the number of PABPN1 foci in non-stimulated neuronal cultures and blocks cLTP-associated increases in small PABPN1 foci, without affecting formation of large foci.

The findings suggest that Arc stabilizes PAPBN1 speckles and specifically regulates formation of small foci as a distinct class. Knockdown of Arc at the single neuron level removes a brake on speckle formation and in so doing also occludes activity-dependent formation. During neuronal activity-dependent gene expression and synaptic plasticity, Arc may provide a homeostatic signal to stabilize PABPN1 speckles.

Given the selective binding of Arc to the PABPN1 RRM, it is interesting to speculate on molecular functions. First, a direct consequence of binding may be inhibition of polyA recognition and polyadenylation of neuronal activity-induced RNAs. Second, Arc may serve as a chaperone for PABPN1 to regulate stability and activity of speckles through liquid-liquid phase-separation. Recent work shows that heat-shock protein 70 modulates formation of TAR DNA-binding protein (TDP-43) condensate through binding to the RRM of RNA-free TDP43 (Yu et al., 2021). Third, in growing oocytes, PABPN1 mediates formation of phase-separated nuclear polyadenylation domains, for the storage and processing of newly transcribed RNA during the maternal to zygotic transition (Dai et al., 2022). Given frequent parallels between germline and neuronal RNA regulation, it is possible that PABPN1 foci regulated by Arc are specialized for storage and processing of neuronal activity-regulated transcripts. As Arc itself interacts with RNA (Pastuzyn et al., 2018; Eriksen et al., 2021), it may recruit activity-induced RNAs to speckles for processing and export. At synapses, Arc is implicated in phase-separation at the postsynaptic density (Hallin et al., 2021; Zhang and Bramham, 2021; Chen et al., 2022) and may have analogous functions in regulation of membrane less nuclear bodies.

The reciprocal coimmunoprecipitation analysis indicates common interactions between Arc, PABPN1, and PSF. PSF is a multifunctional RNA binding protein involved in transcriptional regulation, 3’-end mRNA processing and splicing, and paraspeckle formation, with additional roles in the cytoplasm (Knott et al., 2016; Lim et al., 2020). PSF has been proposed as an active bridge and integrator of nuclear processes (Yarosh et al., 2015). In excitatory glutamatergic neurons, Arc could serve to coordinate PSF and PABPN1 function.

To be able to manipulate the Arc/PABPN1 interaction and establish causal roles, the protein-protein interface needs to be defined. While Arc binds selectively to the PABPN1 RRM, the peptide array analysis identifies potential binding sites for PABPN1 on the Arc coiled coil, N-lobe and C-lobe. We focused on the C-lobe for modeling because of the more extensive binding observed on peptide arrays. However, it is worth noting that complex formation may entail significant conformational changes which were not explicitly modeled in the docking simulations. Furthermore, the assumption was made that it is the PABPN1 dimer that binds to Arc. In principle the results should hold for the monomer binding case as well, since the other, non-interacting subunit of PABPN1 from docking provides only small support for binding through few sparse interactions. Finally, it was not possible to model PABPN1 binding to the coiled-coil region in the full-length Arc protein.

PABPN1 binding to C-lobe peptides occurred along helix-6 to 8, with clusters of critical single residues identified in helix 6 and 8. As illustrated in Figure 5E, this region harbors conserved dimerization motifs from HIV Gag CA that mediate C-lobe/C-lobe binding during capsid formation (Nielsen et al., 2019; Zhang et al., 2019). This region also encompasses a predicted SUMO-interaction motif and Arc has been shown to interact non-covalently with SUMO moieties and undergo enhanced binding during LTP in vivo (Nair et al., 2017). The structural docking model therefore raises the possibility that binding of PABPN1 to the Arc C-lobe could impede Arc oligomerization and interactions with SUMOylated proteins in the nucleus. The alternative models based on AlphaFold suggest more clearly a binding interface between a PABPN1 dimer and the Arc C-lobe, and this unrestrained modeling highlights several of the segments picked up on peptide arrays and alanine scanning. In the absence of experimental structures for the complex, both models can be used further to develop hypotheses and modulate the interaction functionally. One way to screen for interaction surfaces further could employ the recently developed anti-Arc nanobodies against the Arc N-and C-lobe (Ishizuka et al., 2022; Markússon et al., 2022; Godoy Muñoz et al., 2024)

Previous work showed that Arc associates with PML and the acetyltransferase CREB binding protein (CBP) in cultured hippocampal neurons to promote transcriptional downregulation of GluA1 AMPAR subunits and thereby homeostatic downscaling of synaptic inputs to the neuron (Korb et al., 2013). Studies based on ectopic overexpression of Arc in HEK239 cells and hippocampal neurons have demonstrated co-localization of Arc with the histone acetyl-transferase TIP60 along with β-spectrin isoform βSpIVε5 to PML nuclear bodies (Bloomer et al., 2007; Wee et al., 2014), while coimmunoprecipitation shows complex formation between Arc, TIP60, and βSpIVε5. Super-resolution imaging of endogenous Arc and TIP60 in cLTP-treated hippocampal neurons also showed extensive proximity and partial colocalization of the proteins. Functionally, recruitment of TIP60 following ectopic Arc expression is associated with a specific chromatin modification (enhanced H4K12 acetylation) in neurons, suggesting a function in epigenetic regulation (Wee et al., 2014). Notably, studies on the effect of acute cocaine administration indicate that Arc suppresses chromatin remodeling in striatal neurons and regulates behavioral responses to cocaine (Salery et al., 2016). Here, we confirm Arc interaction with TIP60 in adult brain, lending support to previous in vitro studies (Wee et al., 2014; Oey et al., 2015). The major advance of the present work is the identification of endogenous Arc protein complex formation with PABPN1 and PSF as regulators of 3’-end mRNA processing during in vivo synaptic plasticity, with evidence for direct binary interactions between Arc and PABPN1, and Arc-dependent regulation of PABPN1 dynamics. Further work is needed to assess the function of Arc complexes and possible coupling between transcriptional and post-transcriptional regulation.

## Supplemental Information

Supplemental information for online publication is provided in separate files.

## Acknowledgments

Supported by Polish-Norwegian Research Fund (Grant PNFR-96 to GW and CRB); The Research Council of Norway (grants 186115, 204861, 199355, 249951 to C.R.B.),

## Author Contributions

G.W. and C.R.B. conceived the study and obtained funding. T.K. performed the subcellular fractionation, prepared samples for mass spectrometry, conducted all biochemical analysis of Arc interaction complexes, analyzed data and prepared figures. S.P. and F.P. conducted electrophysiological studies for in vivo LTP, dentate gyrus dissection, and performed data analysis and figure preparation. A.S. contributed to immunocytochemical localization of Arc and binding partners in cultured hippocampal neurons. Y.I. contributed to Arc knockdown studies in neuronal cultures. K.P. performed Arc purification for mass spectrometric identification of partners, conducted work on drug-induced seizures, imaging of Arc in brain tissue sections, chemical LTP in cultured neurons, and associated data analysis. D.H-K conducted cLTP experiments and Arc knockdown in hippocampal neurons. A.M. performed confocal microscopy, 3D image analysis, and figure preparation. E.I.H. generated purified GST-fusion proteins in the lab of P.K. and participated in peptide array analysis. J.L and S.D-H. conducted the Haddock structural docking model. P.K. conducted the AlfaFold modeling. C.R.B. supervised the project. T.K., A.M., and CRB wrote the paper, with contributions from all authors.

## Materials availability

This study did not generate new unique reagents.

## Inclusion and Diversity

We support inclusive, diverse, and equitable conduct of research. One or more of the authors of this paper self-identifies as an underrepresented ethnic minority in their field of research or within their geographical location.

