## Supplemental Figure Legends for "Arc/Arg3.1 binds the nuclear polyadenylate-binding protein RRM and regulates neuronal activity-dependent formation of nuclear speckles"

**Figure S1. Immunoblot showing expression of Arc, PABPN1, PSF, SRRM2, and PML protein in DG subcellular fractions**

Immunoblot analysis was performed subcellular fractions (cytosol, membrane, nuclear soluble (nucleosol), nuclear chromatin bound, and cytoskeletal) from HFS-treated and contralateral control DG. Arc and PABPN1 expression was enhanced in the nucleosol of the HFS-treated DG.

**Figure S2. Arc interaction with PABPN1 and PSF persists in RNase treated samples.** The nucleosol fraction from HFS-treated and contralateral DG was RNase treated prior to immunoprecipitation using mouse monoclonal anti-Arc antibody and complexes were separated by SDS-PAGE. The membrane was immunoblotted with anti-PABPN1 and anti-PSF antibodies. Arc interaction with PABPN1 and PSF was detected in control and RNase treated samples. Representative blot from 3 independent experiments.

**Figure S3. Peptide binding array data from full-length protein sequence.** Analysis of full membrane of spotted peptides sequence on cellulose-bound peptide array covering the PABPN1 (A) and Arc (B) sequences with an offset of 7 amino acids, incubated with purified GST-fused Arc or GST-PABPN1, respectively. The areas marked on the sequence are those used for alanine scanning.

**Figure S4. Additional data from Haddock Model.**

(A) Predicted RMSD estimates of the relative positioning of individual domains (chain A, B - PABPN1 dimer, chain C Arc) for 5 predictions with different Alphafold2 models.

(B) Annotation of the human PABPN1 sequence (Uniprot:Q86U42). Yellow highlights mark RNA binding residues according to (Song et al., 2008)

(C) Annotation of the human Arc sequence (Uniprot: Q7LC44)
