## Supplemental Figures for "Arc/Arg3.1 binds the nuclear polyadenylate-binding protein RRM and regulates neuronal activity-dependent formation of nuclear speckles"

Fig. S1

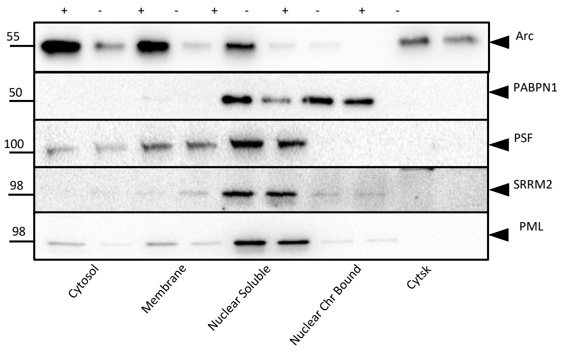


Fig. S2

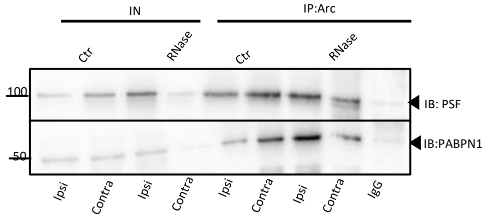

Fig. S3

**A** **Full-length peptide sequence of PABPN1. Area marked in red correspond to the fragment used for alanine scanning**

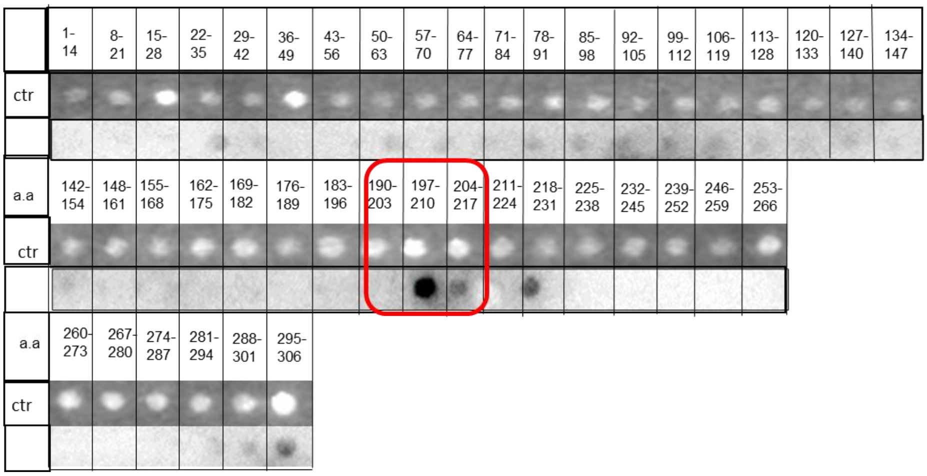


**B** **Full-length peptide sequence of Arc. Area marked in red correspond to the fragment used for alanine scanning**

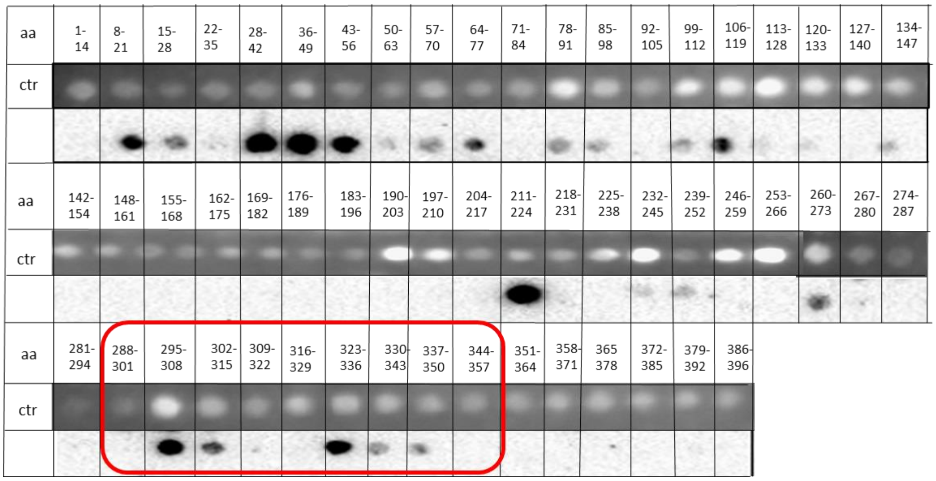

Fig. S4

**A**

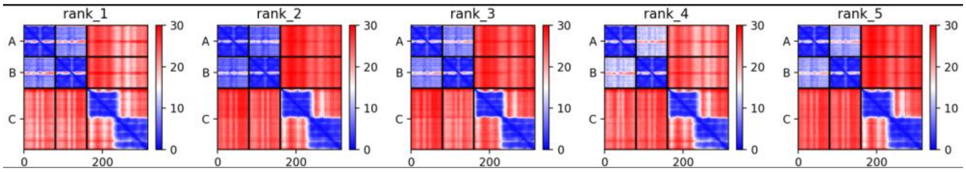


**B**

**polyadenylate-binding protein 2 isoform 1**

UniProtKB - Q86U42 (PABP2_HUMAN) NP_004634

*MAAAAAAAAAAGAAGGRGSGPGRRRHLVPGAGGEAGEGAPGGAGDYGNGLESEELEPEEL*
*LLEPEPEPEPEEEPPRPRAPPGAPGPGPGSGAPGSQEEEEEPGLVEGDPGDG*AIEDPELE
AIKARVREMEEEAEKLKELQNEVEKQMNMS*PPPGNAGPVIM*SIEEKMEADAR**S**I**Y**VGNVD
YGATAEELE**AHFHGCGSVNRVTILCDKFSGHPKGFAY**IEFSDKESVRTSLALDESLF**R**G**R**
QIKVIPKRTNR*PGISTTDRGFPRARYRARTTNYNSSRSRFYSGFNSRPRGRVYRGRARAT*
*SWYSPY*

- predicted disorder
- 3b4m_A,B,C,D – 100% identity structural hit,
- coiled-coil domain PDB:6ytu,
- *HP***K***GF* – lysine residue important for binding according to the peptide array
- **peptide fragments involved in binding according to the peptide array**
- **RNA-binding residues** (according to Song et al, 2008, PNAS)

**C**

***Activity-regulated cytoskeleton-associated protein***

*UniProtKB - Q7LC44 (ARC_HUMAN)*

MELDHRTSGGLHAYPGPRGGQVAKPNVILQIGKCRAEMLEHVRRTHRHLLAEVSKQVERE
LKGLHRSVGKLESNLDGYVPTSDSQRWKKSIKACLCRC**QETIANLERWVKREMHVWREVF**
**YRLERW**ADRLESTGGKYPVGSESARHTVSVGVGGPESYCHEADGYDYTVSPYAITPPPAA
GELPGQEPAEAQQYQPWVPGEDGQPSPGVDTQIFEDPREFLSHLEEYLRQVGGSEEYWLS
QIQNHMNGPAKKWWEFKQGSVKNWVEFKKEFLQYSEGTLSREAIQRELDLPQKQGEPLDQ
F**LWRKRDLYQTLYVDADEEEIIQYVVGTLQPKLKRFLRHPLPKTLEQLIQRGMEVQD**DLE
QAAEPAGPHLPVEDEAETLTPAPNSESVASDRTQPE

- predicted disorder,
- coiled-coil domain,
- **6gse_A** – 97% identity structural hit,
- **peptide fragments involved in binding according to the peptide array**
